# Sniffer beetles: Odor imaging reveals congeneric herbivores identify their congeneric hostplants based on differential olfactory perceptions

**DOI:** 10.1101/2023.01.06.522974

**Authors:** Gauri Binayak, Ashish Deshpande, Neelesh Dahanukar, Prajakta Ingale, Kesavan Subaharan, Vinay Kumar Thirumalahatti Munikrishnappa, Hemant Ghate, Sagar Pandit

## Abstract

Hostplant’ location and conspecific aggregation on the hostplant are the key behaviors of several herbivore insect species. The cues used by insects for host identification and aggregation initiation have been researched mainly using a single hostplant species. The chemical repertoire of plants, including volatile and non-volatile secondary metabolites, is critical in mediating these processes. In natural ecosystems, often several closely related plant species co-occur. Despite these related plant species’ similar chemical repertoires, insects proficiently locate their hosts. How they resolve such complex chemical cues is understudied. To study the basis of such resolution, we used five commonly co-occurring *Ipomoea* spp. as hostplants and four *Chiridopsis* spp. (beetles) as their herbivores. In this wild sympatric system from the Western Ghats of India, monophagous, biphagous, and oligophagous *Chiridopsis spp*. are specialist herbivores of different *Ipomoea* species. We studied the chemistry of these beetles’ stringent host-specificity by determining the roles of chemical cues in hostplant location and aggregation.

We analyzed beetles’ hostplant preferences vis-a-vis hostplant volatile blends. We found plant volatiles as the primary hostplant identification cues. Using GC-MS/-FID and SPME headspace analyses, we characterized odor blends of the five *Ipomoea* spp. and identified putative attractants and repellents for each *Chiridopsis* sp. using multivariate statistics. We determined their attractant or deterrent natures using behavioral assays and ascertained their perception by the antennal olfactory receptors using electroantennography. Beetles responded to these compounds only when they were delivered via their hostplant odor blends. Beetles did not respond when these compounds were given singularly or via non-host odor blends. We infer that these semiochemicals’ attractant, repellent, or neutral characters are associated with the hostplant’s volatile blend-the matrix. We integrated the multi-source data to visualize this in-flight odor perception by representing odor as color variations or ‘odor images.’ Odor imaging revealed beetles’ differential olfactory perception of different hostplants and indicated how a beetle distinguishes between two closely-related plant species. Additionally, it showed a different olfactory perception of the same hostplant by different closely-related beetle species, suggesting they have evolved to recognize the same odor using different components. Our work demonstrates that the hostplant’s odor blend matrix is crucial; beetles do not respond to attractants/ repellents without it. Together, the closely-related plant species form an ideal system to understand how insects perceive subtle differences between hosts and non-host cues in nature. This investigation also underlines the relevance of studying entire odor blends over individual compounds.

## Introduction

Plant-herbivore food webs represent more than 40% of global terrestrial biodiversity, and most of these herbivores are phytophagous insects^1^. The range of hostplants occupied by insect herbivores shows large variability, from one or few plant taxa to several different plant taxa. Hostplant selection by phytophagous insects holds significant importance in an ecological system because this variation among organisms influences several ecological phenomena, such as the coexistence of competitors^2^, persistence of species upon environmental disturbances^3^, and maintenance of inter-species interaction networks^4,5^. Locating and identifying hostplants amid complex mixed vegetation is a challenging but crucial task for insects^6^ and is facilitated by a combination of sensory inputs. Recognizing a plant commences with perceiving olfactory and visual cues from a distance, followed by gustatory and tactile cues that may help host selection after contact^7^. Insects incorporate these sensory inputs into a foraging decision; upon perceiving the correct combination of indicators, a plant is recognized as palatable or unpalatable^6,8^.

Of all these sensory signals, plant volatile organic compounds (VOCs) play a significant role in mediating this interaction as insects often use them to identify hosts and non-hosts while in flight^9–12^. VOCs are used for host identification even by insect herbivore larvae^13–16^, ovipositing females^17–21^, parasitoids^11,22–24^, and even in underground interactions^25^. Research has explored the effects of individual plant VOCs or groups of VOCs on insect behavior. Several studies have successfully identified species-specific plant VOCs that attract or repel their insect herbivores^15,26–32^. On the other hand, other reports suggest that it is the blend of all the released compounds which insects collectively perceive while making a host selection/ location decision^33–36^. To understand hostplant location and identification amidst mixed vegetation, studying plant odor blends rather than single compounds offers a more realistic portrayal of what foraging insects encounter. Plant odor blends are often complex mixtures consisting of hundreds of compounds^12,37,38^. Most identified plant volatiles are ubiquitous across plants as odorants characteristic of plant taxa are rare^12,39–41^, and there is a broad overlap between the odorants that different insects can detect^12,42^. This odor detection overlap within a limited range of plant VOCs suggests that the identification is enabled by the combination of compounds and their signal processing in the insect nervous system. Therefore, it is natural for a foraging insect that the functional unit of plant odor is not a single compound but multiple compounds co-occurring in a blend. There is also evidence that subtle alterations in the proportions of these compounds can drastically affect the host location behavior^43–45^. Further, researchers have demonstrated specific mixtures of volatiles to attract insects when their components did not^46^, and compounds functioning as host cues in a blend have been shown to become non-host cues when presented alone^47^. All these reports support the idea that odor blends have emergent properties and that hostplant identification relies on recognizing the blend rather than individual components^44,47,48^.

Researchers have studied how an insect simultaneously perceives all the components of an odor blend and which features are critical for host recognition. These chemical cues have mainly been studied using a single agricultural pest model or crop species, where large areas of hostplant monocultures are grown^37^. In comparison, insects’ original ecosystems are much more diverse, where they encounter complex odor bouquets from different plant communities, including abundant non-host odor and high background noise^37,44,47,49^. Moreover, such mixed vegetation habitats frequently harbor closely-related co-occurring plant species with similar chemical repertoires. To understand how foraging insects resolve these complex cue mixtures, we must study wild plant-insect systems in natural habitats. In this regard, a wild system of closely related plants and insects can offer valuable insight into understanding host recognition in natural odorscapes. The tortoise beetles and their larvae (Coleoptera: Chrysomelidae), specialist folivores of the plant family Convolvulaceae, offer such a system. Our observations in the northern ranges of the Western Ghats were that among all the genera of these cassidine beetles, species of the genus *Chiridopsis* show exclusive hostplant associations with plants of the genus *Ipomoea*. For example, *C. nigropunctata* is monophagous (feeds on only one *Ipomoea* species), *C. undecimnotata* is biphagous (feeds on two *Ipomoea* species), whereas *C. bistrimaculata* and *C. bipunctata* are oligophagous (feed on more than two *Ipomoea* species). A naturally available negative control in the study system is *I. parasitica*, which co-occurs with the other plant species, but is not fed upon by these beetles. The *Ipomoea*-*Chiridopsis* system is a unique interaction in which the hostplant spectrum of one insect genus is exclusively associated with different species of only one hostplant genus. Even when these congeneric plant and insect species co-occur, we have observed that when beetles are disturbed, they start flying around and return to their hostplant often without landing on the neighboring plants of other species. The *Ipomoea*-*Chiridopsis* system can thus be ideal for understanding host identification and selection cues. Specifically, we asked, upon encountering these closely related plants during foraging, how do these beetles perceive and differentiate the odors of the closely related host and non-host *Ipomoea* plants?

## Materials and methods

### Plants

Seeds of *I. batatas, I. carnea, I. elliptica, I. triloba*, and *I. parasitica* were collected in and around Pune. Plants were grown and maintained in controlled conditions (temperature: 27°C, humidity: 70%, photoperiod: 12 h light and 12 h dark) in a climate chamber. Fresh leaves from grown plants were used for insect behavioral assays, VOC profiling, and maintaining insect cultures in the laboratory. In behavioral assays, fully expanded, healthy, unwounded leaves were used unless otherwise specified.

### Insects

Adults and larvae of *C. nigropunctata, C. undecimnotata, C. bistrimaculata*, and *C. bipunctata* were collected from in and around Pune. Insects were reared on fresh leafy twigs of their host plants in an insectarium with the same controlled conditions as the climate chamber.

### Field observations on natural occurrence

To understand the hostplant preferences of *Chiridopsis* insects, we observed their occurrence on the five *Ipomoea* spp. in their natural habitats of the Western Ghats. Observations were made in three separate sites in three seasons (n= 9). In every site, we counted the total number of ootheca, larvae, and beetles of each species on 10 plants of each *Ipomoea* sp.

### Multiple-choice assays

To understand the feeding preference of *Chiridopsis* beetles in multiple-choice assays, we presented each *Chiridopsis* spp. with fresh leaves of all five *Ipomoea* spp. in an assay jar (height 15 cm, diameter 25 cm). An artificial leaf of average surface area (adaxial+ abaxial) 50± 5 cm^2^ weighing 0.3± 0.05 g (similar to the average surface area and weight of an *I. elliptica* leaf) was cut from Whatman filter paper and included among the choices as a negative control. For each *Chiridopsis* sp., we conducted six assays (n= 6). Considering these insects’ slow feeding rates, we standardized the assay time to 24 h to provide beetles enough time to explore and feed on all given choices with a quantifiable area. We calculated the amount of feeding on each leaf as [(leaf area devoured from a given Ipomoea spp./ total area devoured from all leaves) × 100].

### No-choice assays

To understand the feeding behavior in the absence of the most preferred hostplant, we conducted no-choice assays for each beetle species, where one individual was exposed to a fresh leaf of a single *Ipomoea* sp. at a time. Each assay consisted of 5 cages with a single *Ipomoea* sp. for 6 h (n= 30). We calculated the feeding amount on each *Ipomoea* sp. as surface area devoured (mean± SE).

### Survivorship assays

To study the survivorship of insects on the different hosts, we released 20 individuals of each *Chiridopsis* sp. (per plant) on each *Ipomoea* sp. (n= 5). Plants were caged to prevent insects from escaping. Insects were allowed to feed on the plants for five days. We counted the survivor number on each plant in each assay and calculated the survivorship as [(number of survivors on a given *Ipomoea* sp./ 20) × 100] (mean± SE).

### Total odor blend complementation assay

Volatile organic compounds were extracted from the fully expanded, healthy unwounded leaves of *I. batatas, I. carnea, I. elliptica, I. triloba*, and *I. parasitica* by solvent extraction. In a screw-cap glass vial with silicone septa, 5 mL dichloromethane (DCM) was used to extract volatiles from 1 g of leaf tissue for 2 h. Extracts were dehydrated using anhydrous sodium sulfate (Rankem, India) and further concentrated to 1 mL using a vacuum concentrator (Labconco, Kansas City, MO, USA). Concentrated extracts were incubated overnight at −80°C to precipitate high molecular weight lipids, which were then removed by centrifugation at 10000 rpm for 10 min at 4°C. Extracts were further concentrated to a final volume of 250 μL and stored in air-tight glass autosampler vials (Chromatography Research Supplies, India) at - 20°C till further use. These DCM extracts majorly contain plant odorants; therefore, we will henceforth refer to the *Ipomoea* leaf extracts as ‘odor blends.’ We complemented five artificial leaves, each with the odor blend of a single *Ipomoea* sp. (physiological concentration). Every *Chiridopsis* spp. was subjected to multiple-choice assays between the five odor-complemented leaves (n= 20). A DCM-complemented and a non-complemented artificial leaf were included in each assay as controls. Beetle visits on each artificial leaf were quantified to understand whether plant odor is the primary hostplant identification cue.

### Extraction and analysis of *Ipomoea* odor blends

We extracted VOCs from *I. batatas, I. carnea, I. elliptica, I. triloba*, and *I. parasitica* leaves by the solvent extraction method described earlier. As an internal standard for quantification, we spiked the extraction solvent DCM with nonyl acetate (2.2 μg/ mL). Compounds were identified and quantified using a gas chromatograph (7890B GC system, Agilent Technologies) coupled to a mass spectrometer and flame ionization detector (7000D GC/triple quadrupole and FID, Agilent Technologies, Santa Clara, CA, USA). Compounds were separated on a DB-5MS capillary column (30 m × 0.32 mm i.d. × 0.25 μm film thickness, Agilent Technologies) using helium as carrier gas with a flow rate of 2 mL/ min. Column temperatures were programmed as follows: 40 °C hold time 5 min, ramp 1: 5 °C/ min till 180 °C, ramp 2: 20 °C/ min till 280 °C, hold time 5 min. Mass spectra were obtained using 70 eV electron ionization with a scan time of 0.2 s for m/z 30-600. Compounds were identified using mass spectral libraries NIST11 and Wiley (8th edition). Compounds’ Kovat’s retention indices were calculated using an n-alkane ladder (C7-C21). Compounds were quantified on GC-FID, where concentrations of different volatiles were normalized with nonyl acetate^50^.

### Statistical analyses

Quantitative data (number of insects on different *Ipomoea* spp., feeding preference, and survivorship on different *Ipomoea* spp.) were analyzed by one-way analysis of variance (ANOVA), and the statistical significance (*p*≤ 0.05) was determined by Fisher’s least significant difference on StatView software (ver. 5.0).

We performed a principal component analysis (PCA) to understand how the five *Ipomoea* spp. differ in their VOCs. PCA was performed on a correlation matrix to account for scale differences in various compounds. To understand whether the plant spp. form significantly different clusters with respect to their VOCs we performed an analysis of similarity (ANOSIM) using Manhattan distances to account for high dimensionality in the data. ANOSIM was performed with 9999 permutations at two levels, at the level of all groups together, with the null hypothesis that all groups are the same and pairwise comparison between groups. When multiple pairwise tests were performed, we used sequential Bonferroni correction to account for family-wise errors. Analysis was performed using PAST 3.26.

We performed a partial least square (PLS) analysis to understand the correlation between multiple dependent variables (feeding preferences of *Chiridopsis* spp.) and multiple independent variables (plant volatiles). The null hypothesis was that there was no correction between the given variables and the first and second axes of PLS, and this was tested using a one-sample t-test. We performed multiple regression between the feeding preferences of *Chiridopsis* spp. and *Ipomoea* volatile compounds to understand whether the plant VOCs determine the observed feeding preferences. The null hypothesis that the standardized coefficient of the multiple regression was not significantly different from zero was tested using multiple one-sample t-tests. We used sequential Bonferroni correction to account for family-wise errors. Analysis was performed using XLSTAT®.

### Complementation assays

To understand the function of compounds correlated with *Chiridopsis* hostplant identification, we conducted complementation assays with all candidate compounds commercially available as analytical standards.

#### Complementation of individual compounds on artificial leaves

For each *Chiridopsis* spp., each putative attractant and repellent was complemented on an artificial leaf in serially increasing concentrations (n= 20 for each concentration). The first concentration pasted was equal to half the physiological concentration in the most-preferred host. The next used concentration was equal to the physiological concentration, doubling the compound’s concentration in the test leaf (a 2-fold increase). Further increments were 4, 6, 8, and 10 folds. In addition to assays using the compounds correlated with beetle preference, we also performed assays using the following technical controls:

- Neutral compound: compound detected in *Ipomoea* spp. but not correlated to beetles’ preferences (β-caryophyllene)
- Foreign compounds: Compounds not detected in any of the five *Ipomoea* spp. Three foreign compounds were used: hexanal (a green leaf volatile), *(Z)*-3-nonen-1-ol (an aliphatic alcohol), and valencene (a sesquiterpene).

At each increment, we recorded beetles’ preferences using a dual choice assay (1 h duration), including compound-complemented (test) and solvent-complemented (control) leaf choices. To estimate attraction towards or deterrence from the test leaves, we calculated the percentage of beetles preferring test and control leaves on their first visits (mean± SE).

#### Complementation of individual compounds on host and non-host leaves

To find how these compounds operate within a blend, we serially raised the concentration of each attractant and repellent in two most-preferred plants and two least-preferred plants of each *Chiridopsis* sp. (n= 10 per concentration). This was done by exogenously applying the compound over the leaf in increasing concentrations, thereby raising its total concentration in the test leaf to 1.5, 2, 4, 6, 8, and 10 folds. At each increment, we observed beetles’ preference for 1 h in a dual choice assay between a test and control leaf. We calculated preference for a choice as the percentage of beetles who visited and initiated feeding on it (mean± SE).

#### Complementation of individual compounds on artificial leaves along with host and non-host odor blends

We also applied attractants and repellents in increasing concentrations as described earlier (n= 10 per concentration) on artificial leaves along with odor blends of the most-preferred plants or the non-host *I. parasitica*. In these assays, the filter paper artificial leaves used were cut in the shape and size of the *Ipomoea* leaf in consideration. At each increment, we observed beetles’ preference for 1 h in a dual choice assay between a test and control leaf. Preference for a choice was calculated as the percentage of beetles who visited and initiated feeding on it (mean± SE).

### Solid phase microextraction (SPME) headspace analysis

We ascertained the presence of the candidate compounds in the *Ipomoea* headspace by SPME using a fiber assembly (divinylbenzene/ carboxen/ polydimethylsiloxane, needle size 24 ga; Sigma, India). All SPME fibers were conditioned before use by inserting them into the gas chromatograph injector (270 ºC, 1 h) as provided in the instruction manual. For each species, a potted plant was enclosed in a ventilated glass cylinder and exposed to an SPME fiber for 1 h to collect headspace volatiles. Soil was covered with a polypropylene bag to minimize the release of soil volatiles into the headspace. The same setup without a plant was used as a blank. For analysis of collected headspace volatiles, the fiber was inserted in the injector of the gas chromatograph and identified using mass spectrometry as described earlier.

### Electroantennography

The electrophysiological response of beetle antennae to the candidate compounds was tested using an electrographic system (Syntech, Hilversum, The Netherlands) consisting of a dual electrode probe for antenna fixation, a CS-05 stimulus controller and an IDAC 232 box for data acquisition. Each antenna was fixed between the two electrodes using Spectra 360 conductive gel (Parker, Orange, New Jersey) as described by Reinecke *et al*.^51^. The antenna was continuously flushed with a stream of activated charcoal-filtered air. Solutions of authentic standards were prepared in DCM, and 10 μL from each was applied to a filter paper strip. The solvent was allowed to evaporate before placing the strip in the apparatus. A purified airstream (pulse time 0.5 s, continuous flow 25 ml/ s, pulse flow 21 ml/ s) flowing over the antennal preparation delivered the stimulus puff. A time delay of 20 s was maintained between consecutive stimulus puffs. The antennal responses were recorded as voltage deflections (mV) through a high-impedance probe connected to an amplifier (IDAC-4, Syntech). The blank stimulus was DCM. We conducted 10 replicates for six concentrations (0.625, 1.25, 2.5, 5, 10, and 20 ppm) of all compounds. Each EAG response was corrected for solvent and background effects by subtracting the response to DCM from the response to the stimulus.

### Odor imaging

We generated odor images for every beetle-plant pair to visualize olfactory perceptions as color variations. For this, we multiplied the concentration of each attractant, repellent, and neutral compound with the beetle’s standardized regression coefficient for that compound. The resulting values were plotted as a square pie diagram for every beetle-plant pair. Leaf shapes were pixelated using the pie charts such that each pixel was a pie chart.

## Results

### Occurrence of *Chiridopsis* spp. on *Ipomoea* spp. shows a distinct preference pattern in nature

We observed that the four beetle species occur only on certain *Ipomoea* spp. in natural habitats (Fig. 1A), exhibiting a distinct pattern. *C. nigropunctata* was found only on *I. elliptica* (Fig. 1B-D), and *C. undecimnotata* was found only on *I. elliptica* (majorly) and *I. batatas* (Fig. 1E-G). On the other hand, *we found C. bistrimaculata* and *C. bipunctata* on four hostplants-*I. batatas, I. carnea, I. elliptica*, and *I. triloba*, displaying a preference for some over others (Fig. 1H-M). We did not observe any ootheca, larvae, or beetles on *I. parasitica*. The hostplant-specific occurrence of each *Chiridopsis* sp. showed the same trend at various developmental stages, such as ootheca, larvae, and adults.

**Figure 1:**
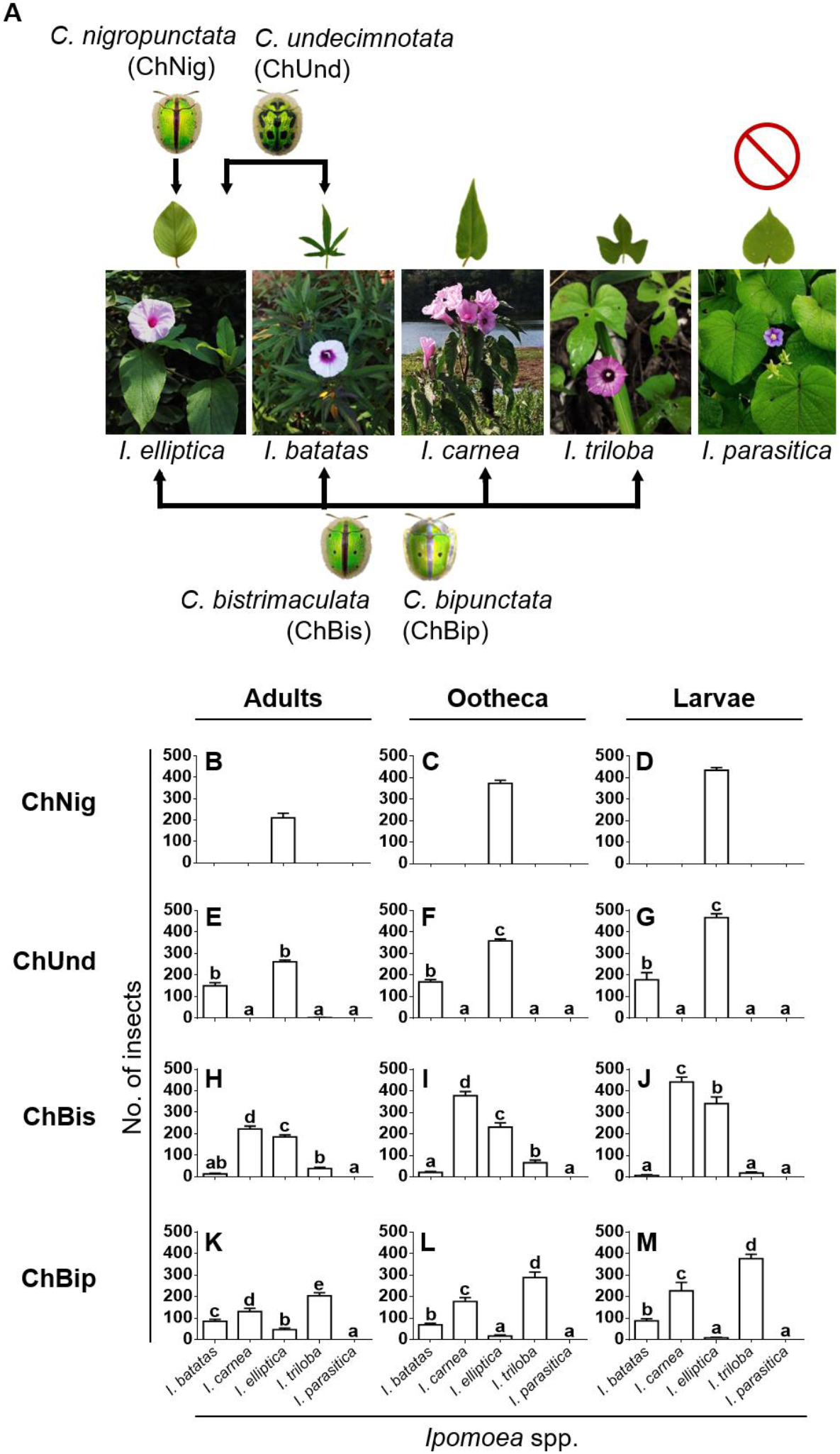
Natural occurrence of *Chiridopsis* spp. on *Ipomoea* spp. (**A**) Arrows indicate occurrence. Beetle names have been given abbreviations that will be used throughout the figures. Observations from the wild are that there is a high specificity in the co-occurrence of each *Chiridopsis* spp. with specific *Ipomoea* spp. No *Chiridopsis* spp. has been found on *I. parasitica*. Occurrence of adults, ootheca, and larvae of (**B**) to (**D**) *C. nigropunctata*, (**E**) to (**G**) *C. undecimnotata*, (**H**) to (**J**) *C. bistrimaculata* and (**K**) to (**M**) *C. bipunctata* was measured on different *Ipomoea* plants in their natural habitats. We counted insect numbers in 3 field locations in 3 seasons (n= 9). In each location, the number of insects on each *Ipomoea* sp. was considered as the total on ten individuals. Data is plotted as mean± SE. Hostplant preferences of each species followed the same trend in every stage of its life cycle. Different letters denote significant differences (*p*≤ 0.05, one-way ANOVA).

### *Chiridopsis* spp. maintain stringent hostplant preferences under controlled conditions

We conducted choice and no-choice assays under controlled laboratory conditions to experimentally validate our field observations. When presented with multiple choices simultaneously, feeding preferences of the *Chiridopsis* spp. (percentage area fed on each leaf in the assay) closely followed the trend of their natural occurrences (Fig. 2A-D). *C. nigropunctata* was strictly monophagous on *I. elliptica* (Fig. 2A), whereas *C. undecimnotata* fed only on *I. elliptica* and *I. batatas*, preferring the former nearly five times more than the latter (Fig. 2B). *C. bistrimaculata* and *C. bipunctata* were oligophagous on *I. elliptica, I. batatas, I. carnea*, and *I. triloba* (Fig. 2C, D). However, they exhibited some preferences: *I. carnea* was most preferred, and *I. batatas* was least preferred by *C. bistrimaculata*. Contrarily, *C. bipunctata* most preferred *I. batatas* and least preferred *I. carnea*. In all assays, no beetle fed on *I. parasitica*. We observed the same trend in egg-laying choice assays (Fig. S1). We also conducted no-choice assays to observe whether these patterns alter upon the unavailability of the preferred plants. In almost all cases, *Chiridopsis* beetles maintained their hostplant range and showed similar preferences as described above (Fig. 2E-H). *C. nigropunctata* remained monophagous on *I. elliptica* and strictly avoided all other plants. When provided its two hostplants separately, *C. undecimnotata* equally fed on both, not displaying a preference for *I. elliptica* over *I. batatas*. Although scantily, it also fed on *I. triloba* (a non-host in nature). The oligophagous *C. bistrimaculata* fed on all hostplants without a preference for one. *C. bipunctata*, however, maintained *I. batatas* and *I. carnea* as its most preferred and least preferred host, respectively. *I. parasitica* remained as a non-host.

**Figure 2:**
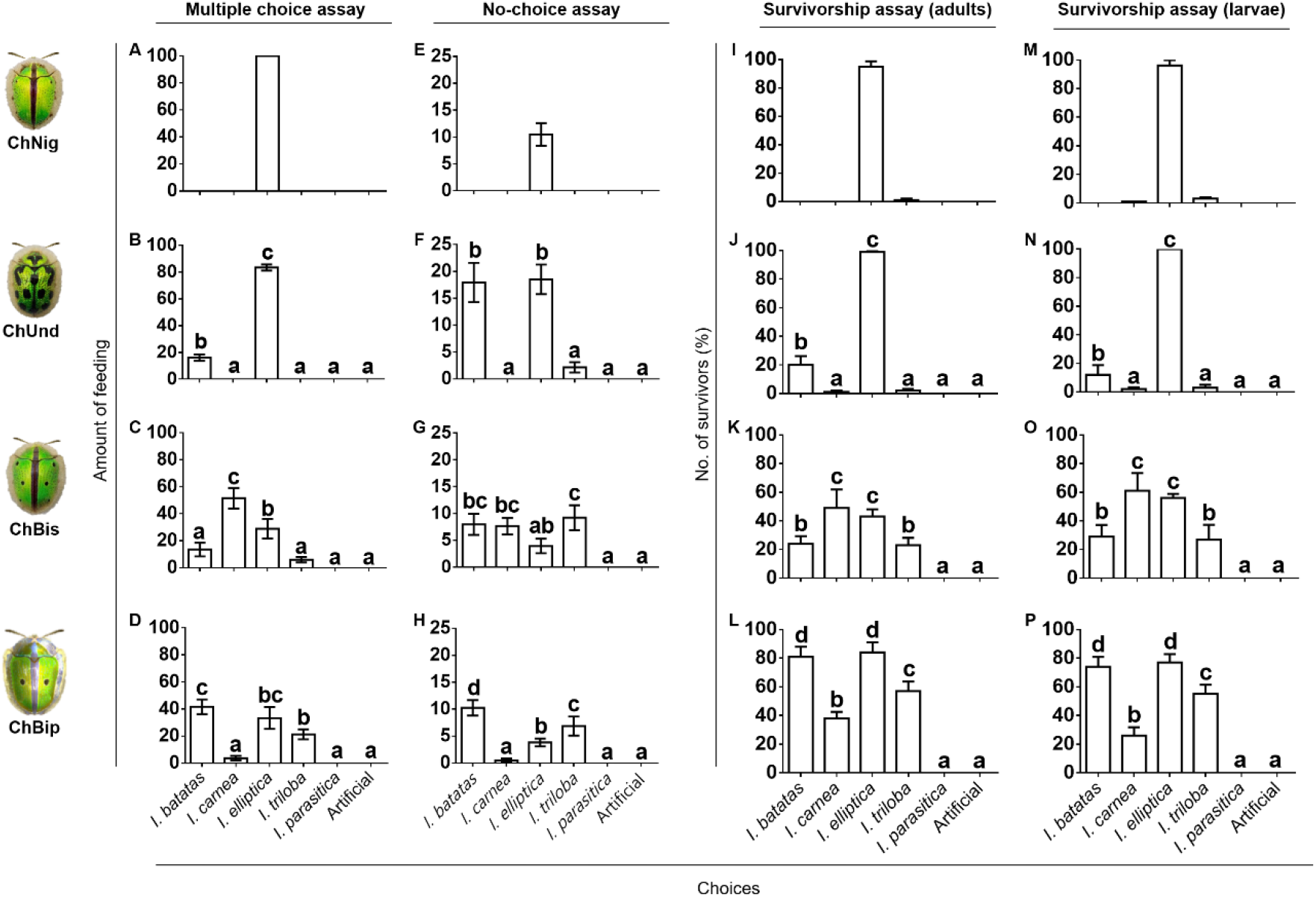
*Chiridopsis* spp. show highly stringent hostplant preferences in feeding. (**A**) to (**D**) Multiple choice assays established the mono-, bi-, and oligophagous nature of the *Chiridopsis* spp. Hostplant preferences were measured as percentage area (mm^2^) fed on each *Ipomoea* sp. leaf in 24 h (mean± SE, n= 6). (**E**) to (**H**) No-choice assays confirmed that the hostplant associations of the *Chiridopsis* spp. do not change even in the absence of the most preferred hostplants. The amount of feeding was measured as the area (mm^2^) fed on each *Ipomoea* sp. leaf (mean± SE, n= 30). Feeding preferences of each *Chiridopsis* spp. shows the same trend as their survivorship on the different plants, in both adult (**I**) to (**L**) and larval (**M**) to (**P**) life stages (n= 5 assays, each containing 20 insects). Different letters in graphs denote significant differences (*p*≤ 0.05, one-way ANOVA).

### Insects’ hostplant preferences follow the trend of their survivorship

In survivorship assays, we observed the percentage of larval and beetle survivors to be highest on their respective most-preferred hosts and lowest on least-preferred hosts (Fig. 2I-P). *C. nigropunctata* and *C. undecimnotata* had 100% survivorship on their most preferred hostplant, *I. elliptica* (Fig. 2I, J, M, N), on which they fed heavily. On non-hosts, most *C. nigropunctata* and *C. undecimnotata* insects refrained from feeding and died of starvation. Concurrent with the above trends, *C. bistrimaculata* and *C. bipunctata* fed and survived on all *Ipomoea* spp. except *I. parasitica* (Fig. 2K, L, O, P). For these oligophagous insects, 100% survivorship was not observed on any hostplant. Despite this, the survivorship on different plants closely resembled the trend of their feeding choice. Survivorship of *C. bistrimaculata* was the highest on its most preferred host, *I. carnea*, and lowest on its least preferred host, *I. batatas*. Survivorship of *C. bipunctata* was highest on its most preferred host, *I. batatas*, and lowest on its least preferred host, *I. carnea*. We also observed 100% mortality in all insects exposed to *I. parasitica*, with no trace of feeding on any plants.

### Hostplant odor is the major host identification cue

When each *Chiridopsis* sp. was simultaneously subjected to the odor extracts of the five *Ipomoea* spp. (Fig. 3A), beetles paid more visits to artificial leaves complemented with their hostplants’ odor than non-complemented ones (Fig. 3B-E). The trend closely followed the trend of feeding preferences, with highest visits to the most preferred odor, and fewer visits to the least preferred hostplant odor. Beetles paid no visits to non-host odors. The observation that hostplant odor alone was sufficient to prompt beetle visits suggests that host identification for the *Chiridopsis* spp. is strongly associated with olfactory cues.

**Figure 3:**
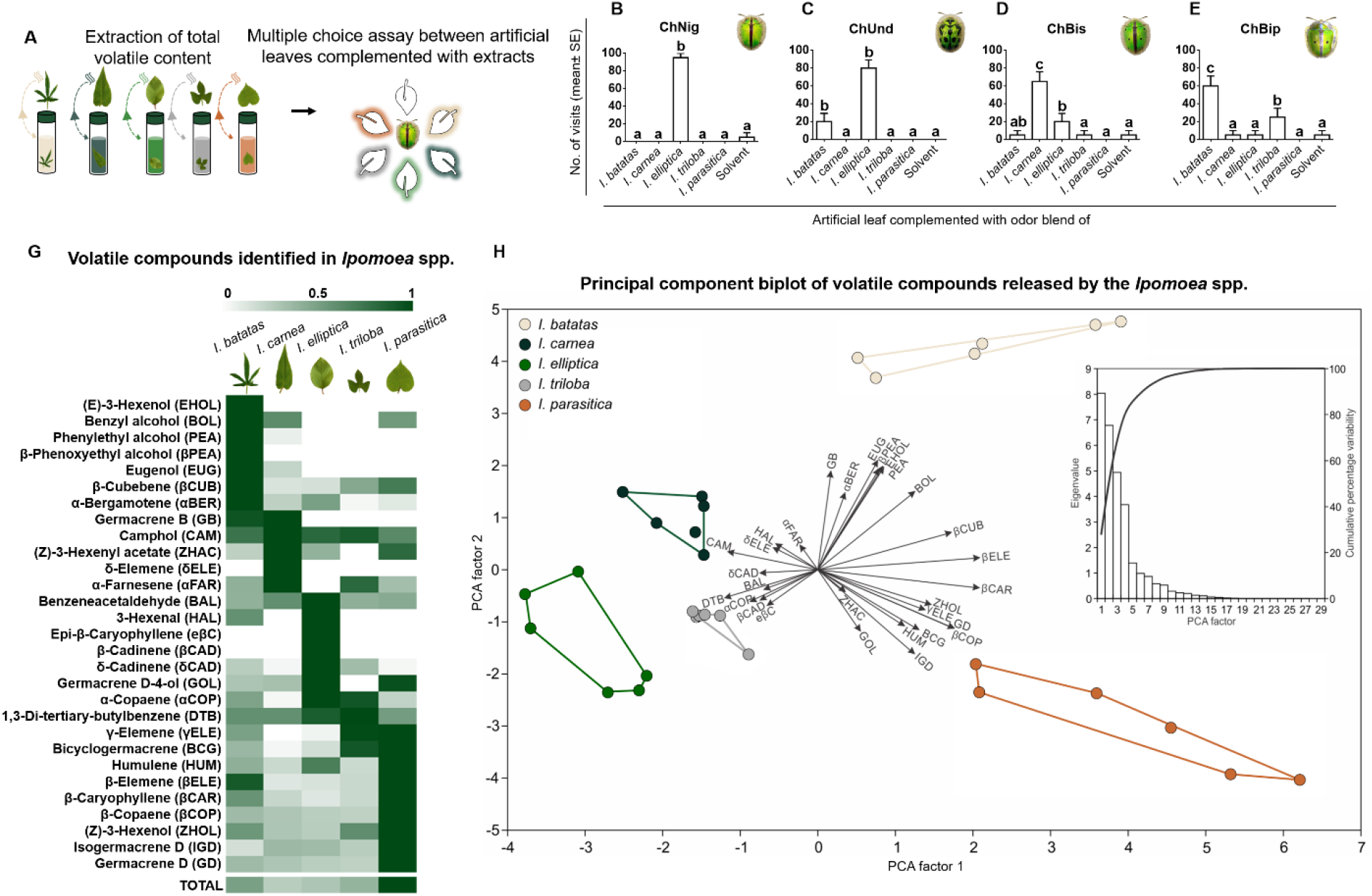
The five *Ipomoea* spp. release characteristic odor blends that are necessary and sufficient to cause beetle visits. (**A**) Schematic of odor blend complementation assay. Total volatile blend was extracted from each *Ipomoea* spp. and pasted on artificial leaves. Each beetle’s preference was assayed when simultaneously provided artificial leaves complemented with different odor blends. An artificial leaf coated with solvent was used as a control. (**B**) to (**E**) Number of visits to each plant’s blend closely resembles the trend of feeding preferences. Different letters denote significant differences (*p*≤ 0.05, one-way ANOVA, n= 20). (**C**) The different *Ipomoea* spp. are associated with distinct compositions of volatile compounds. Columns in the heatmap represent *Ipomoea* spp., and rows represent compounds. Each cell represents the mean concentration of a compound (nmol/ g) relative to nonyl acetate internal standard. Values in each row are normalized to the highest value in that row (n= 6). All compounds have been given abbreviations that will be used throughout subsequent figures. (**D**) Principal component biplot of factor scores of plant species and factor loadings of volatiles. A scree plot of eigenvalue and percentage variation explained by each PCA factor is provided in the inset. Pairwise comparisons of clusters using ANOSIM based on Manhattan distances showed that all clusters are significantly different.

### *Ipomoea* spp. produce characteristic odor blends

Leaf volatile organic compounds (VOCs) of the five *Ipomoea* spp. were extracted and analyzed by gas chromatography-mass spectrometry and gas chromatography-flame ionization detection (GC-MS and GC-FID). We identified 29 compounds, each present in unique combinations and concentrations in each *Ipomoea* sp., thereby generating five signature odors (Fig. 3G, Table S1). Sesquiterpenes contributed to most of the odor for all species, (90.8-95.9%). The quantitative contribution of aldehydes (0.2-3.8%), oxygenated terpenes (0.02-1%), and other compounds (0.5-1.9%) to the blends was relatively minor. Total volatile content was highest in the herbivore-resistant *I. parasitica* (3655.13 ± 194.37 nmol/ g leaf). The sesquiterpene germacrene-D was the most quantitatively dominant compound in all five odor blends. *I. parasitica* contained ≥ 3.9-fold higher concentration (1433.6± 145.45 nmol/ g) than the other *Ipomoea* spp. (Table S1). We performed a principal component analysis (PCA) and analysis of similarity (ANOSIM) using Manhattan distances to understand how these five odor compositions compare. PCA separated the five *Ipomoea* species based on their VOCs on the first two axes (Fig. 3H). On the first PCA axis, *I. batatas* and *I. parasitica* separated from *I. carnea, I. elliptica*, and *I. triloba* based on high factor loading for volatile compounds like *(Z)*-hex-3-en-1-ol, γ-elemene, β-cubebene, β-elemene, β-caryophyllene, β-copaene, germacrene-D, and bicyclogermacrene, and low factor loading for compounds like camphol and 1,3-ditertiary-butylbenzene (Table S2). Clusters of plant species with respect to their VOC composition were significantly different (ANOSIM, R= 0.7565, *p*= 0.0001).

### Hostplant preferences of *Chiridopsis* spp. are associated with specific plant VOCs

PLS analysis helped visualize the relationships between the 29 VOCs, five *Ipomoea* spp., and four *Chiridopsis* spp. by plotting them as vectors. The angle between two vectors indicates their relation to each other (Fig. 4A). The monophagous *C. nigropunctata* and biphagous *C. undecimnotata* showed similar VOCs associated with their feeding behavior. Both their preferences strongly correlated to the first component of PLS, which also positively correlated with compounds such as β-cadinene, 3-hexenal, epi-β-caryophyllene, benzeneacetaldehyde, and α-copaene, and negatively correlated with β-caryophyllene, α-farnesene, γ-elemene, benzyl alcohol, β-elemene and (*Z*)-hex-3-en-1-ol (Fig. 4A, Table S3). The similarity in plant VOCs they correlate to could explain their common preference of *I. elliptica* as a hostplant.

**Figure 4:**
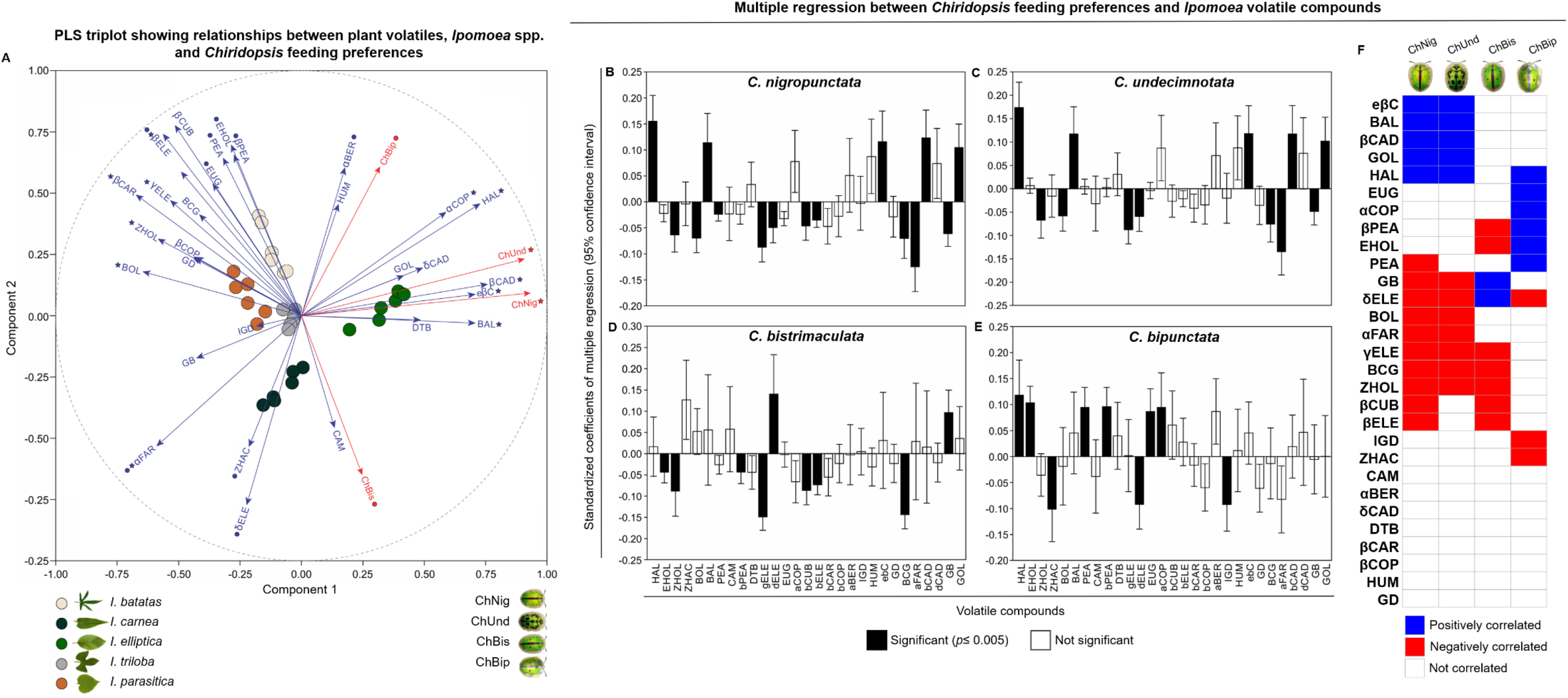
Host preference of each *Chiridopsis* spp. correlates with certain volatile compounds in *Ipomoea* odor blends. (**A**) Partial least squares (PLS) regression triplot of factor scores of *Ipomoea* species, factor loadings of volatiles and factor loadings of *Chiridopsis* feeding preference as estimated by multiple choice assays (Fig 2A-D). Independent volatiles are shown as blue vectors, while dependent variables are shown as red vectors. Correlations are absolute on the dashed unit circle. Filled star with the name of the variable indicates that the variable is significantly correlated on the first PLS axis, filled circles indicate that the variable is significantly correlated on the second PLS axis, while both stars and circles indicate that variables are significantly correlated on both the axes. Significance is assessed after sequential Bonferroni correction (Table S3). (**B-E**) Standardized coefficients of multiple regression between feeding preference and plant volatiles. Dependent variable feeding preference of (**B**) *C. nigropunctata*, (**C**) *C. undecimnotata*, (**D**) *C. bistrimaculata* and (**E**) *C. bipunctata*. Error bars are 95% confidence intervals. Bars in black are significant after sequential Bonferroni correction (*p*≤ 0.005) (Table S5). (**F**) Summary of PLS regression results showing volatile compounds that show positive (blue), negative (red) and no (white) correlations with *Chiridopsis* feeding preferences on various *Ipomoea* spp.

Although *C. bistrimaculata* and *C. bipunctata* have the same hostplants, their feeding preferences correlated oppositely. *C. bistrimaculata* negatively correlated, whereas *C. bipunctata* positively correlated with the second component of PLS, to which α-bergamotene, β-phenoxyethyl alcohol, (*E*)-hex-3-en-1-ol, phenylethyl alcohol, eugenol, β-cubebene, and β-elemene positively correlated, and δ-elemene, α-farnesene, and (*Z*)-3-hexenyl acetate negatively correlated (Fig. 4A, Table S3). Therefore, despite sharing a host range, these two beetles associate with different VOCs, resulting in opposite relationships.

Multiple regression between the *Ipomoea* VOCs and *Chiridopsis* feeding preferences and analysis of standardized coefficients showed several significant relationships (Fig. 4B-E, Table S4, Table S5). *C. nigropunctata* and *C. undecimnotata* significantly correlated with the same VOCs; positively with 3-hexenal, benzeneacetaldehyde and β-cadinene, epi-β-caryophyllene, and germacrene-D-4-ol, and negatively with (*Z*)-3-hex-3-en-1-ol, benzyl alcohol, γ-elemene, δ-elemene, bicyclogermacrene, α-farnesene and germacrene-B (Fig. 4B, C, F). Additionally, *C. nigropunctata* negatively correlated with phenylethyl alcohol, β-cubebene, and β-elemene, while *C. undecimnotata* did not. *C. bistrimaculata* and *C. bipunctata* correlated oppositely to some volatiles, such as δ-elemene, (*E*)-3-hex-3-en-1-ol and β-phenoxyethyl alcohol (Fig. 4D, E, F). For each *Chiridopsis* spp., we categorized the positively and negatively correlated plant volatiles as putative attractants and repellents, respectively.

### *Chiridopsis* spp. respond to attractant and repellent volatiles only when presented within a hostplant’s odor

We experimentally tested the candidate volatiles’ functions (putative attractants and repellents) through a series of complementation assays (Fig. 5, S2, S3, S4). Presenting individual candidate compounds in serially increasing concentrations neither attracted nor deterred beetles (Fig. S2A, C, E, G). On the contrary, when their concentration was serially increased on each beetle’s two most-preferred plants and two least-preferred plants, the compounds were indeed of attractant or repellent nature as predicted by the multiple regression (Fig. 5). At each concentration of an attractant on a hostplant, most beetles preferred the complemented leaf. Moreover, with increasing concentrations of pasted attractant, complemented leaves were preferred by an increasing number of beetles. Conversely, when we complemented repellents on hostplants, most beetles preferred the non-complemented leaf. As we increased the repellent concentration, fewer beetles preferred the complemented hostplant leaf (Fig. 5A-D). We observed this attraction/ deterrence behavior only when we pasted attractants/ repellents on the beetles’ natural hostplants. On non-host leaves, beetles’ behavior remained unaffected. The beetles did not exhibit this increased attraction/ deterrence upon increasing concentration of compounds that did not correlate with host preference (neutral) or compounds that had not been detected in the *Ipomoea* spp. (foreign) (Fig. S3A-D), thus empirically validating the multiple regression results. These observations also held when the candidate compounds were pasted on artificial leaves pre-coated with hostplant odor blends (Fig. S4A-H), showing that the behavioral responses observed were due to olfactory signals alone.

**Figure 5:**
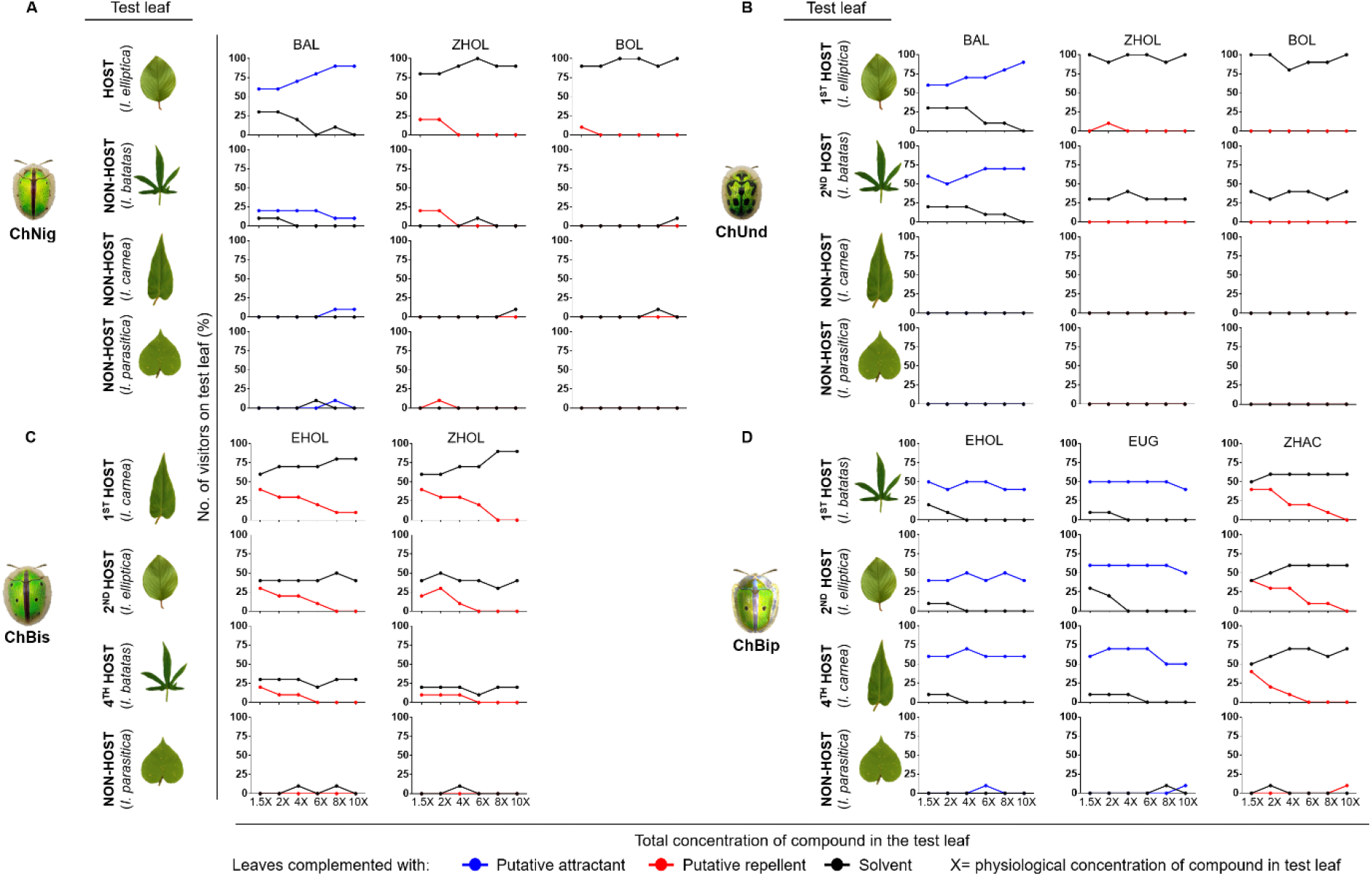
*Chiridopsis* spp. respond to attractant and repellent volatiles only within a hostplant’s odor blend. On the leaves of each *Chiridopsis* sp.’s two most preferred and two least preferred plants, we incrementally raised the concentration of putative attractants and repellents. Beetles were subjected to dual choice assays between leaves pasted with a putative attractant/ repellent (test) and leaves pasted with only solvent (control). Preference of (**A**) *C. nigropunctata*, (**B**) *C. undecimnotata*, (**C**) *C. bistrimaculata*, and (**D**) *C. bipunctata* were analyzed by quantifying the number of beetles who visited each choice. The figure shows the mean percentage of visitors on control and test leaves (n= 10). Contrary to when we presented these compounds individually (Fig S2), all beetles exhibited behavioral attraction or deterrence when the compounds were presented along with leaves naturally releasing their respective odor. As each attractant/repellent’s concentration was incremented, more/ fewer beetles visited the leaf. Notably, this behavioral response was displayed only when the test leaf was of a natural host. When the test leaf was a non-host, increasing levels of attractants/ repellents did not result in respectively more/ fewer visits. In addition to putative attractants and repellents, as technical controls, we also serially raised the concentration of a compound not correlated with beetle feeding (neutral compound) and compounds not detected in the *Ipomoea* spp. (foreign compounds). In these cases, no preference was shown by beetles (Fig S3). Together these results indicate that these attractants and repellents exert their function only when presented within a hostplant’s odor blend.

### Candidate attractants, repellents, and neutrals are present in the headspace and are EAG-active

Detection of the experimentally tested VOCs in the *Ipomoea* headspace (Fig. S5) by SPME-HS analysis ascertained that plants release the compounds into the headspace. EAG analysis showed that all the attractant, repellent, and neutral test compounds elicited an electrophysiological response in beetles’ antennae, suggesting that they are olfactorily perceived by receptors. Increased responses were observed with a corresponding increase in stimulus concentration, with saturation beyond a threshold concentration in some cases (Fig. S6).

### Odor imaging

The proportions of attractants, repellents, and neutrals in each *Ipomoea* sp. were plotted as a pie diagram for all the *Chiridopsis* sp. (Fig. S7). Odor images generated from these proportions for all beetle and plant combinations revealed that the beetles perceive the five plant odors differently, visualized as color differences (Fig. 6). For all beetles, the first host appeared with an attractant blue hue due to the higher proportion of attractants (Fig. 6C, H, L, P, R). The hues of green and red were seen in odor images of less-preferred hostplants or non-hosts due to their higher proportions of neutral and repellent compounds. The odor images also reveal that different beetles perceive the same hostplant using different attractants, repellents, and neutrals and thus perceive the same plant with different odor images (Fig. 6A-P, B-Q, C-R, D-S, E-T).

**Figure 6:**
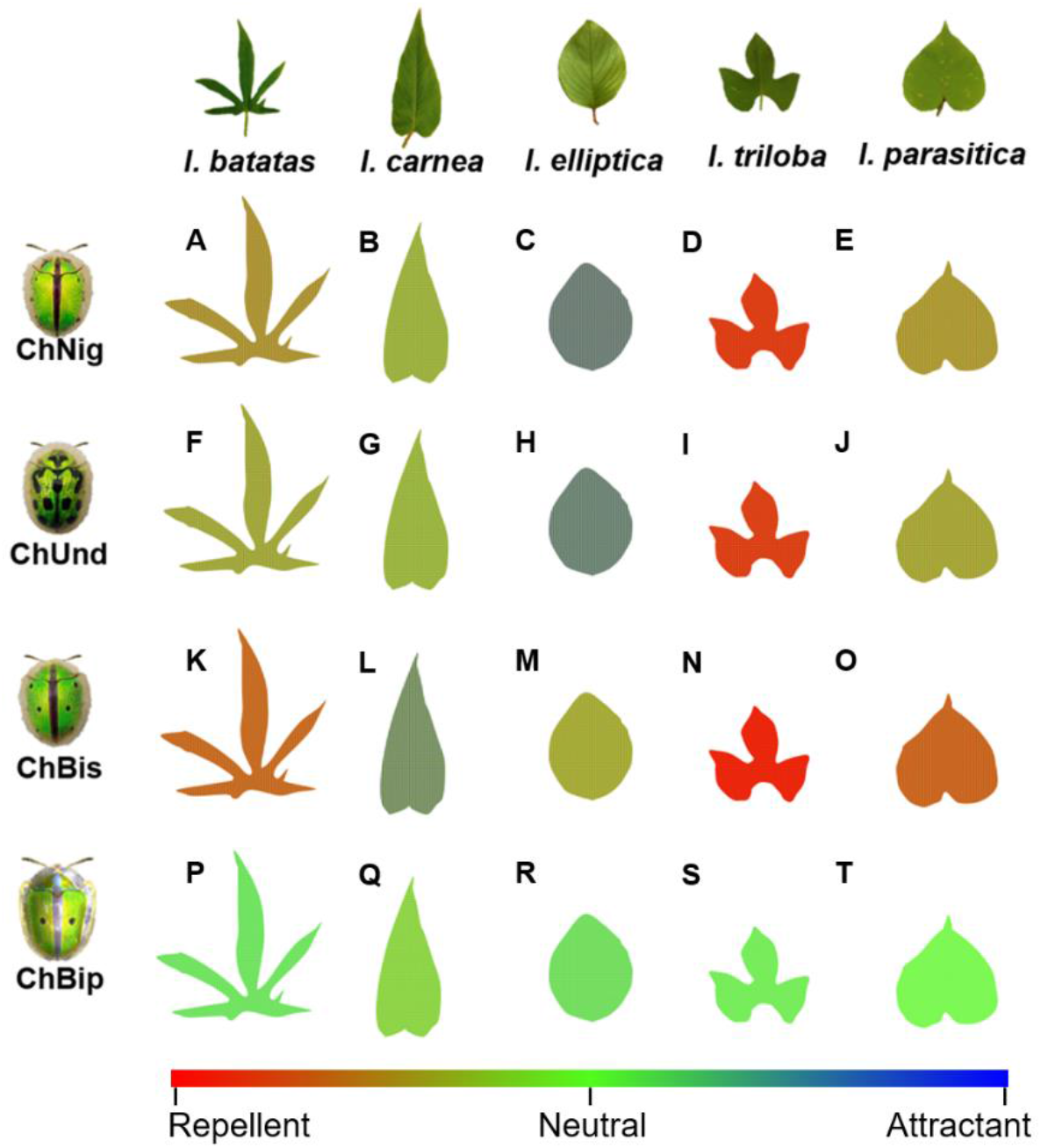
Each *Ipomoea* spp. is associated with a characteristic odor image for each *Chiridopsis* spp. For every beetle-plant pair, the pie diagrams in Fig. S7 were used to pixelate respective leaf shapes, thereby causing all 20 leaves to gain different hues. The odor images show how each beetle perceives the five *Ipomoea* spp. differently. Leaves of most-preferred hosts appear with a blue hue due to a higher proportion of attractants (**C, H, L, P, R**), whereas those of less-preferred hosts (**F, K, N, Q, S**) or non-hosts (**A, B, D, E, G, I, J, O, T**) appear with a yellow or red hue due to higher proportion of neutrals or repellents. When odors are similar, their resolution is based on the proportions of attractants and repellents. The odor images also show how the same plant is perceived with different odor images by different beetle species, depending on their preferences. Through this novel odor imaging tool, we attempted to visually represent beetles’ in-flight perceptions of host and non-host odors.

## Discussion

In the *Ipomoea-Chiridopsis* interaction, the host preference spectrum of one insect genus is exclusively associated with different species of only one genus of hostplants. In this naturally sympatric system, the insects show highly host-specific occurrences, which they maintain through every life cycle stage. Consistent with these field observations, we observed a striking trend in the hostplant preferences displayed by these beetles in our laboratory experiments. The choice of hostplants to feed displayed the same trend as the adult and larval survivorship on the different plants. While we observed 100% survivorship of *C. nigropunctata* and *C. undecimnotata* on their most-preferred hosts, *C. bistrimaculata* and *C. bipunctata* displayed survivorship on all their hosts without 100% survival on any one host. This was not surprising as oligophagous insects often tend to have moderate viability on each host and no maximal fitness on a single host^52–54^. Together our results indicate a fine-tuned relationship between these beetles and plants, suggesting that the insect choices are evolutionary adaptations for greater survival rather than spontaneous foraging decisions.

The high specialization observed in this insect-hostplant system, especially considering that all species co-occur, led us to investigate the basis of the precise host identification. Insects are known to begin the hostplant identification process in-flight, using contactless visual and olfactory cues. Despite variable weather conditions and the low resolution of insect vision, we have observed that these beetles land on their hosts directly, suggesting that visual cues are not the major signals associated with *Chiridopsis* host location. On the other hand, the role of olfactory signals in hostplant location by insects is well recognized^6,55,56^. Our observation that hostplant odor alone was sufficient to elicit beetle visits indicated that the major host identification signal in this system is plant odor. GC-MS-FID-based profiling revealed that the *Ipomoea* species are associated with a similar set of VOCs, but the five blends differ in their proportions and concentrations of these compounds. PCA and ANOSIM analysis showed that the five congeneric sympatric plants had significantly different odors due to these quantitative differences. Understanding how the 29 plant VOCs in five *Ipomoea* spp. correlate with hostplant preferences of four *Chiridopsis* spp. required a multivariate statistical approach to deal with the high dimensionality of the data. PLS analysis and multiple regression helped visualize the relationships between these dimensions and revealed that some VOCs are associated with the feeding preferences of each beetle. In this regard, a high resemblance exists between monophagous *C. nigropunctata* and biphagous *C. undecimnotata*. Both beetles positively and negatively correlate similarly with a group of VOCs; this explains their common choice of *I. elliptica* as the most-preferred hostplant. The only correlated VOCs differentiating these beetles (phenylethyl alcohol, β-cubebene, and β-elemene) are higher in *I. batatas* than *I. elliptica*, thus explaining their negative correlation to *C. nigropunctata*, for who *I. batatas* is a non-host. Interestingly, despite sharing the same range of hostplants, oligophagous *C. bistrimaculata* and *C. bipunctata* correlate negatively and show nearly opposite trends of preferences within their four *Ipomoea* hosts. Compounds that are attractants for one are repellents for the other, and vice-versa. Their host preferences correlate with different VOCs, suggesting that these two sympatric beetles recognize the same plants using different cues.

Through a series of complementation assays, we demonstrated how the correlated compounds function as attractants or repellents, and our results agree with statistical correlations. The observation that beetles do not respond to putative attractants and repellents when presented singly, whereas they do respond to odor blends, led us to hypothesize that the attractant and repellent volatiles are functional only when they co-occur with other hostplant volatiles in a blend. Complementation assays supported our hypothesis and demonstrated that these compounds are critically associated with the background volatiles-the matrix. Compounds attracted and repelled beetles only when their levels increased within the hostplant-they did not have the same behavioral effect when increased in non-hosts. Furthermore, these assays showed that the attractive nature of an attractant is not brought about by the specific concentration found in a beetle’s most-preferred hostplant. Similarly, for a repellent, it is not the specific concentration found in the non-host *I. parasitica* which renders it deterrent. If this were the case, increasing attractants on non-hosts would have attracted beetles, and increasing repellents on hosts would have deterred them. Instead, we demonstrate that rather than the absolute concentration of every attractant or repellent, its co-occurrence with other hostplant compounds in a matrix confers it attractant or repellent nature. Therefore, these signals are functional only with the background of all the other compounds, together forming the plant’s odor.

Together, our experiments led us to discover that *Chiridopsis* beetles’ odor blend perception is contextual: the same compound can be of attractant, repellent, or neutral nature, depending on the odor background. Our finding agrees with several other reported host identification studies, which found that background odor critically affects odor perception in insects^47,57,58^. The presence of other hostplant volatiles, even if repellent by themselves, has been found to make some compounds more attractive to insects and aid in host recognition^59^. This could be because such non-attractant, non-host, or repellent compounds function as habitat cues, providing an essential context to the insect that the attractants have indeed originated from a hostplant in nature. In some other cases, the presence of other volatiles has been found to have masking or distracting effects, making key attractant compounds less effective^60^.

The general principles underlying the collective perception of attractants and repellents for hostplant identification have remained poorly understood^60^. We attempted to visualize this olfactory perception by odor imaging. Two components form an *Ipomoea* sp.’s odor image: the composition of VOCs in that plant and how each VOC affects beetle preference. While the first component distinguishes the five *Ipomoea* spp. from each other, the second distinguishes how the four *Chiridopsis* spp. perceive the same *Ipomoea* sp. Each odor image is a visual representation of a particular beetle’s olfactory perception of a particular plant. Together, they suggested that the concentration of attractants and repellents is instrumental within the host odor blend. If two odors are similar, their differentiation is based on the proportions of attractants and repellents. Our use of multiple insect herbivores sharing a hostplant range provided additional insight, as it showed that the olfactory cues are beetle-specific; the identity of attractant, repellent, and neutral compounds in the same hostplant differs for each beetle. As a result, the same plant is perceived as a different odor image by different beetle species, who have evolved different behavioral responses upon perception of the same odorant from a shared host. The integration of behavior, statistics, and metabolomics to image odor perception is the first effort of its kind in the field. Odor imaging sheds light on how a flying beetle perceives its hostplant odor and distinguishes between odor blends of closely related plant species, especially when these plants occur together. These results indicate that olfactory cues are one of the principal factors associated with hostplant recognition and specialization. This tool could be fine-tuned in the future by incorporating odor detection thresholds, volatility, and VOC emission rates.

Host identification by odor perception has been studied over decades in several insect systems. Studies suggest that insects use individual compounds (attractant or repellent) or their mixtures in specific ratios to identify hostplants. Our findings are more in agreement with the latter. Through our study, we demonstrate that VOCs are not independent components to study in isolation; instead, more multivariate studies on entire blends are required to understand how they may function collectively. We have attempted such a multidimensional approach to understanding plant odorscapes, and this is the first study of this kind. Such studies could provide insights into this research area as it offers a realistic perspective on understanding host/ non-host recognition by a foraging insect.

## Supporting information

Binayak et al_Supplementary information

## Acknowledgments

Authors thank Ministry of Human Resource Development, India, for financial support and funding the GC-MS/FID system, Mr. Greg D. Silva, IISER Pune, for inputs, and Mr. Keerthi Raj, IISER Pune, for photography. GB thanks the Council of Scientific and Industrial Research, India, for Ph.D. fellowship; AD thanks SERB, India, for the N-PDF fellowship; authors thank IISER Pune for providing the *on-campus* field site, Mr. G. Pingalkar for field support, and Mr. G. Pawar for help in field experiments. Authors thank Maharashtra State Biodiversity Board for permission to access biological resources. KS and VM thank ICAR NBAIR, Bangalore, for EAG facility.

## Author contributions

GB, AD, ND, KS, and SP conceived and designed the experiments. All authors performed the experiments and collected the data. ND and GB conducted the statistical analyses. All authors interpreted and discussed the results. GB and SP wrote the manuscript with inputs from all the authors; SP acquired funds, administered the project, and supervised the research.

## The following supporting information is available for this article

**Fig. S1** *Chiridopsis* spp. show hostplant specificity in egg-laying.

**Fig. S2** *Chiridopsis* spp. do not respond to volatile compounds presented individually.

**Fig. S3** *Chiridopsis* spp. do not respond to volatiles that are uncorrelated to their feeding preferences or those not detected in the *Ipomoea* spp.

**Fig. S4** *Chiridopsis* spp. respond to attractant and repellent volatiles only within a hostplant’s odor blend.

**Fig. S5** Experimentally verified attractants, repellents, and neutrals are present in *Ipomoea* headspace.

**Fig. S6** Experimentally tested attractants, repellents and neutrals are EAG-active.

**Fig. S7** Each *Ipomoea* sp.’s odor blend has a signature proportion of attractants, repellents, and neutrals, which differs for every *Chiridopsis* sp.

**Table S1** Volatile compounds of *Ipomoea* spp.

**Table S2** Factor loading, percentage contribution, and squared cosines of variables (volatile compounds) on the first two principal components of PCA.

**Table S3** Correlation coefficients and *p* values for volatile compounds (independent variables) and *Chiridopsis* feeding preference (dependent variables) on the first two principal components.

**Table S4** Intercept (*β*_*0*_) and coefficients (*β*_*i*_) of multiple regression (*y* = *β*_0_ + ∑ *β*_*i*_*x*_*i*_) between feeding preference (*y*) and volatile metabolites (*x*_*i*_).

**Table S5** Standardized coefficients and *p* values for the null hypothesis that coefficients are not significantly different from zero.

