## Supplementary material for "Sniffer beetles: Odor imaging reveals congeneric herbivores identify their congeneric hostplants based on differential olfactory perceptions": Binayak et al_Supplementary information

**The following supporting information is available for this article:**

**Fig. S1** *Chiridopsis* spp. show hostplant specificity in egg-laying.

**Fig. S2** *Chiridopsis* spp. do not respond to volatile compounds presented individually.

**Fig. S3** *Chiridopsis* spp. do not respond to volatiles that are uncorrelated to their feeding preferences or those not detected in the *Ipomoea* spp.

**Fig. S4** *Chiridopsis* spp. respond to attractant and repellent volatiles only within a hostplant's odor blend.

**Fig. S5** Experimentally verified attractants, repellents, and neutrals are present in *Ipomoea* headspace.

**Fig. S6** Experimentally tested attractants, repellents and neutrals are EAG-active.

**Fig. S7** Each *Ipomoea* sp.'s odor blend has a signature proportion of attractants, repellents, and neutrals, which differs for every *Chiridopsis* sp.

**Table S1** Volatile compounds of *Ipomoea* spp.

**Table S2** Factor loading, percentage contribution, and squared cosines of variables (volatile compounds) on the first two principal components of PCA.

**Table S3** Correlation coefficients and *p* values for volatile compounds (independent variables) and *Chiridopsis* feeding preference (dependent variables) on the first two principal components.

**Table S4** Intercept ($\beta_0$) and coefficients ($\beta_i$) of multiple regression ($y = \beta_0 + \sum \beta_i x_i$) between feeding preference ($y$) and volatile metabolites ($x_i$).

**Table S5** Standardized coefficients and $p$ values for the null hypothesis that coefficients are not significantly different from zero.

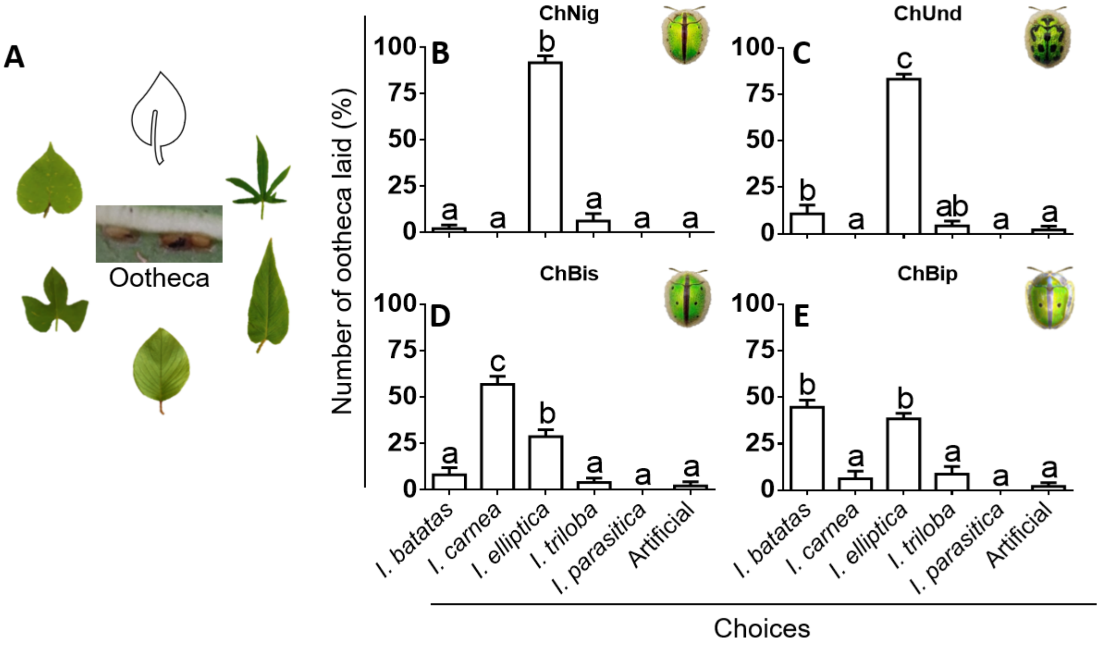


**Figure S1: *Chiridopsis* spp. show hostplant specificity in oviposition.** (**A**) Schematic of oviposition choice assays. Oviposition preference of (**B**) *C. nigropunctata*, (**C**) *C. undecimnotata*, (**D**) *C. bistrimaculata,* and (**E**) *C. bipunctata* were measured as the percentage of ootheca laid on each *Ipomoea* spp. (mean± SE). Different letters denote significant differences ($p \leq 0.05$, one-way ANOVA, n= 5).

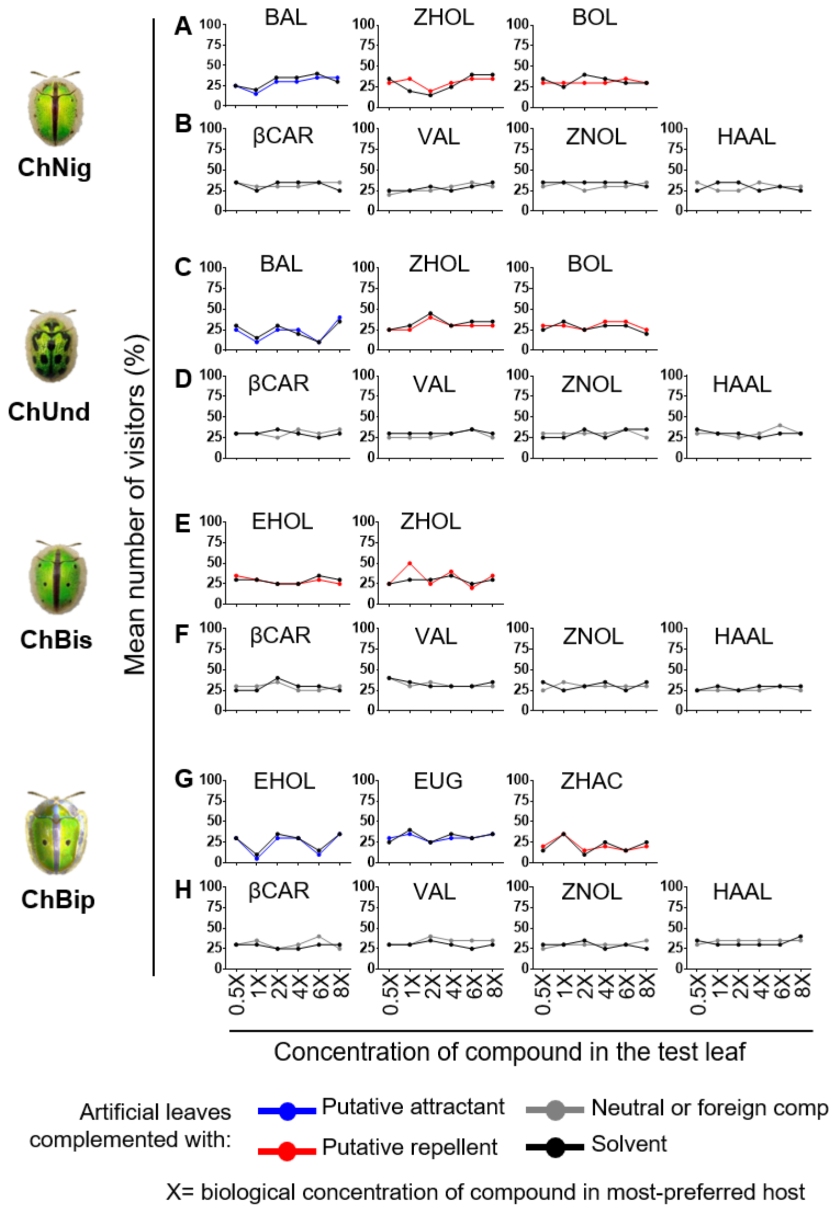


**Figure S2: *Chiridopsis* spp. do not respond to volatile compounds presented individually.** Putative attractants and putative repellents (**A, C, E, G**) for *C. nigropunctata*, *C. undecimnotata, C. bistrimaculata,* and *C. bipunctata* were complemented individually on artificial leaves in increasing concentrations. In addition to putative attractants and repellents, we also performed assays using the following technical controls: compound detected in *Ipomoea* spp. but not correlated to beetles' preferences (neutral compound: β-Caryophyllene) and compound not detected in any of the five *Ipomoea* spp. in the study (foreign compounds: hexanal, (*Z*)-3-nonen-1-ol, and valencene) (**B, D, F, H**). Since these compounds were not detected in the *Ipomoea* spp., the reported physiological concentration in their close relatives was considered as 1X (see Materials and Methods section). Beetles were subjected to a dual choice assay between compound-complemented artificial leaves and control (solvent-complemented) leaves. Behavioral response to the test compounds was estimated as the number of visitors to each leaf. The figure shows the mean percentage of visitors on control and test leaves (n= 20). In all cases, beetles visited both control and complemented leaves similarly, showing no significant preference.

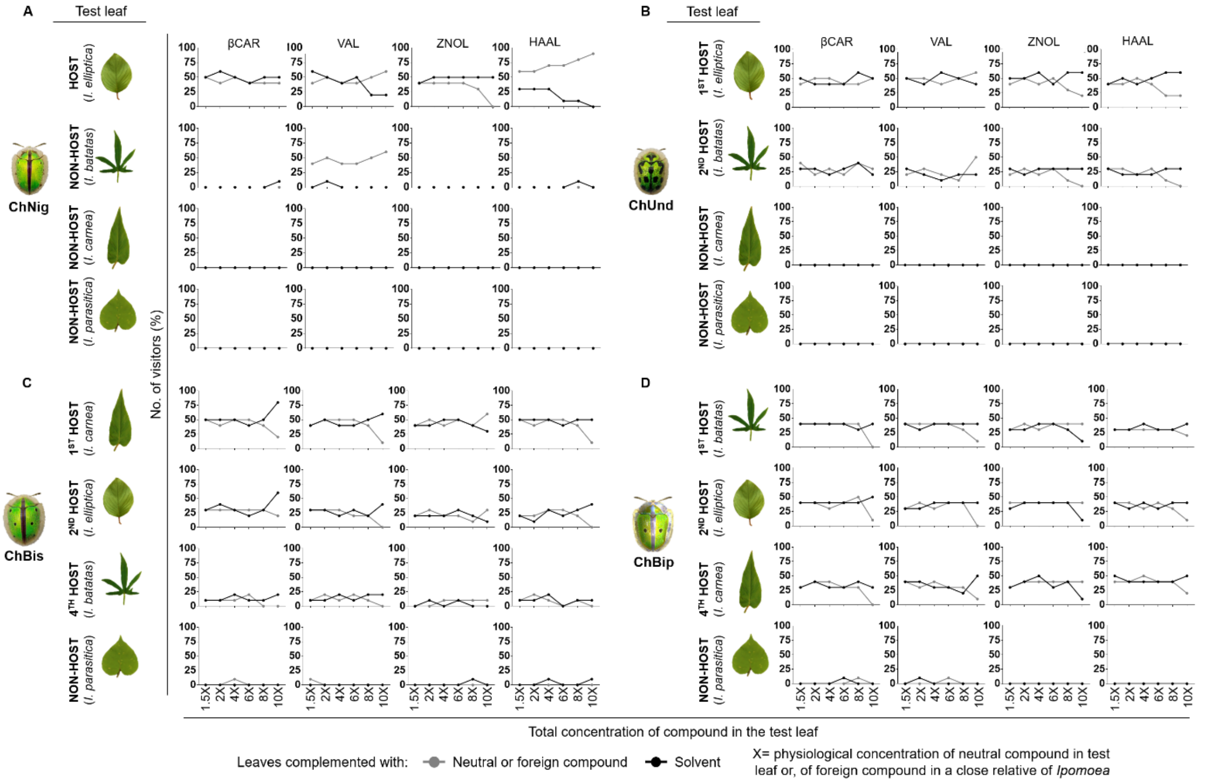


**Figure S3: *Chiridopsis* spp. do not respond to volatiles that are uncorrelated to their feeding preferences or those not detected in the *Ipomoea* spp.** In addition to putative attractants and repellents (Fig 5), as technical controls we also serially raised the concentration of a compound not correlated with beetle feeding (neutral compound: β-caryophyllene) and compounds not detected in the *Ipomoea* spp. (foreign compounds: valencene, (*Z*)-3-nonen-1-ol, and hexanal). Since these compounds were not detected in the *Ipomoea* spp., the reported physiological concentration in their close relatives was considered as 1X (see Materials and Methods section). Beetles were subjected to dual choice assays between leaves pasted with compound (test) and leaves pasted with solvent (control). Preference of (**A**) *C. nigropunctata*, (**B**) *C. undecimnotata*, (**C**) *C. bistrimaculata,* and (**D**) *C. bipunctata* were analyzed by quantifying the number of beetles who visited each choice (n= 10). Data shown in the figure is percentage of visitors on the choices (mean± SE).

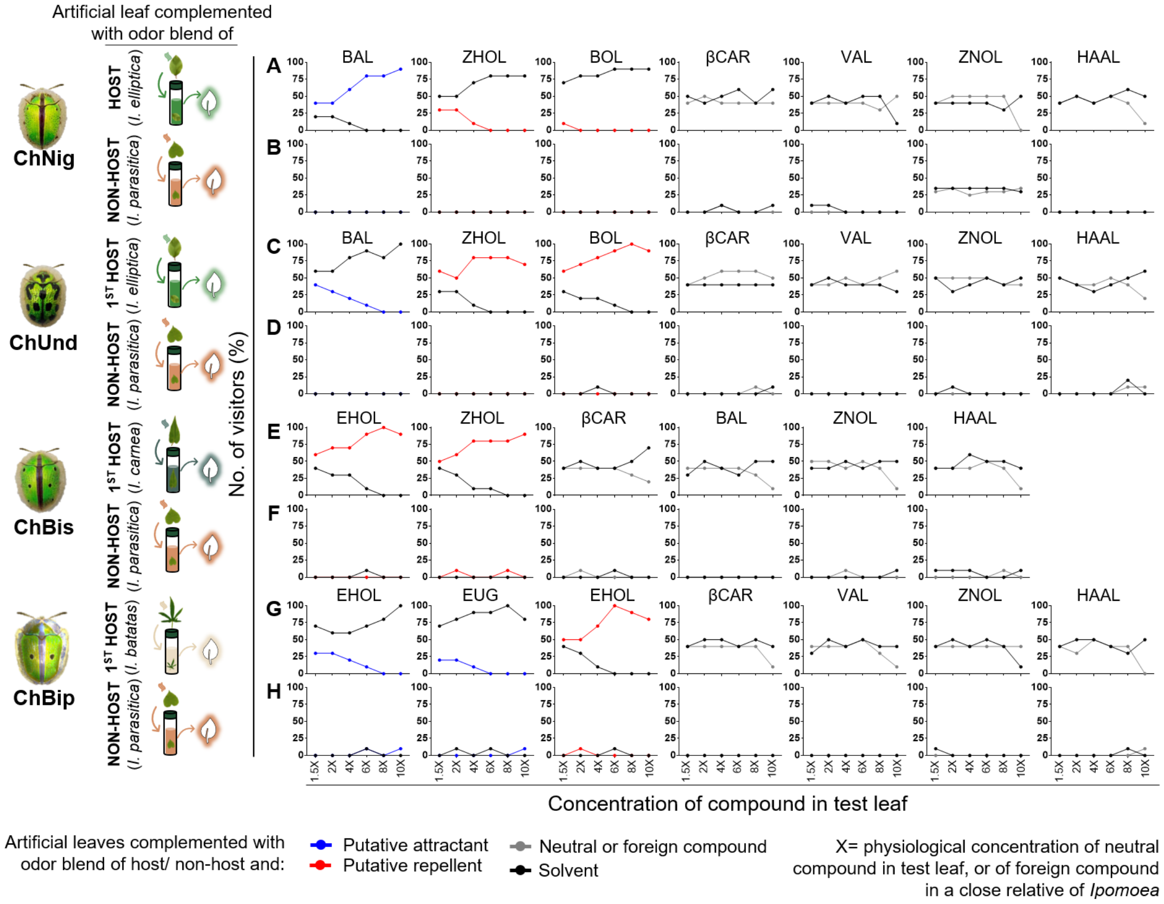


**Figure S4:** ***Chiridopsis*** **spp. respond to attractant and repellent volatiles only within a hostplant's odor blend.** On artificial leaves already coated with the odor blends of most-preferred hosts or non-hosts, we serially increased the concentrations of putative attractants and repellents for each *Chiridopsis* sp. Beetles were subjected to dual choice assays between artificial leaves coated with only odor blend (control) and those coated with a blend+ putative attractant or repellent (test). Preference of (**A, B**) *C. nigropunctata*, (**C, D**) *C. undecimnotata*, (**E, F**) *C. bistrimaculata,* and (**G, H**) *C. bipunctata* were analyzed by quantifying the number of beetles who visited each choice. The figure shows the mean percentage of visitors on control and test leaves (n= 10). Similar to when these compounds were encountered on different leaves (Fig 5), all beetles exhibited behavioral attraction or avoidance when these compounds were encountered within odor blends. Moreover, as the concentration of attractant/ repellent was incremented, more/fewer beetles visited the leaf. This behavioral response was displayed only when the background odor on the test leaf belonged to a natural host. If the pasted blend was of a non-host, increasing levels of attractants/ repellents did not result in more/fewer visits. Together these results indicate that these attractants and repellents exert their function only when present within a hostplant's odor blend.

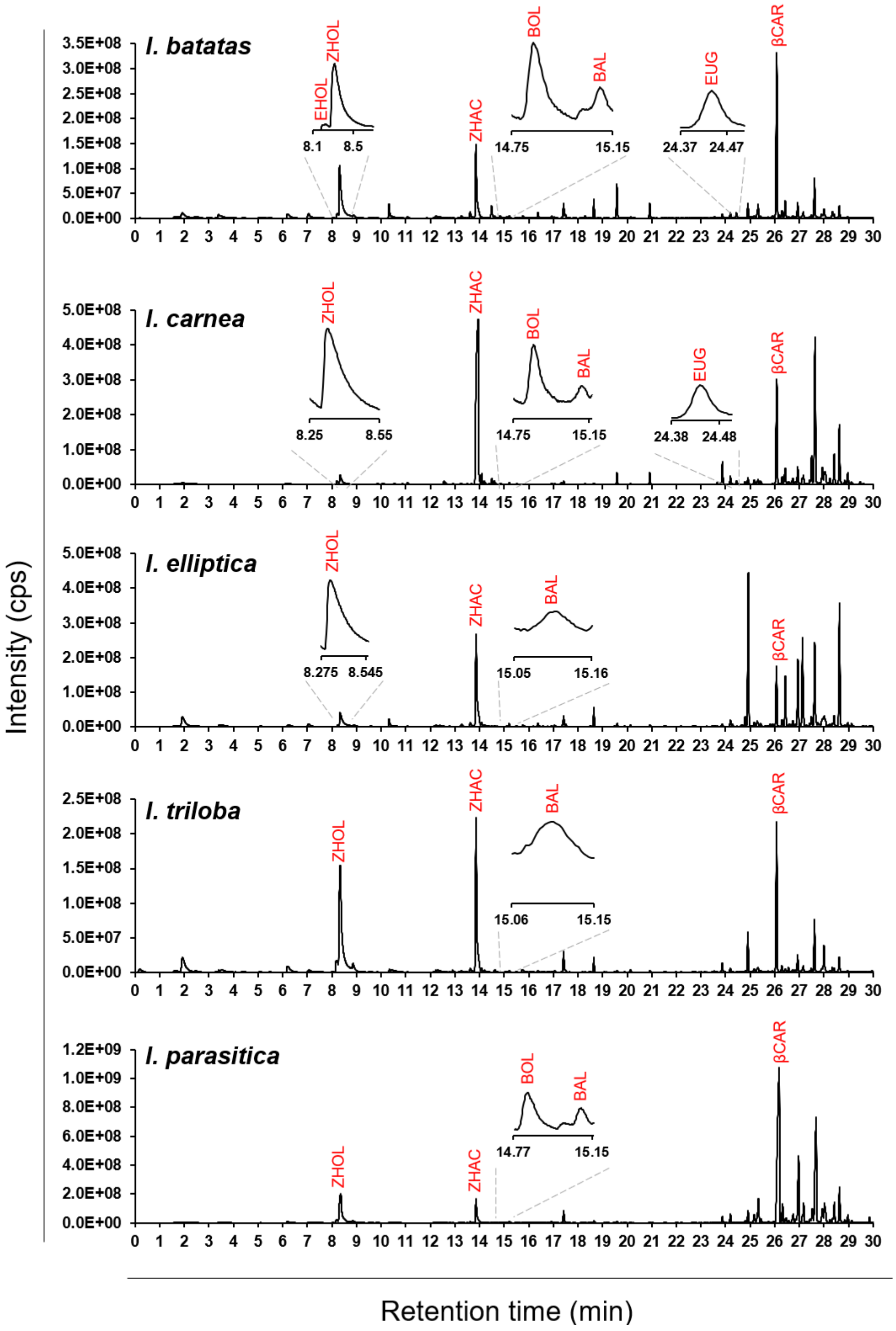


**Figure S5: Experimentally verified attractants, repellents, and neutrals are present in *Ipomoea* headspace.** For each species, a potted plant was enclosed in a ventilated glass cylinder and exposed to an SPME fiber assembly (divinylbenzene/ carboxen/ polydimethylsiloxane) for 1 h to collect headspace volatiles. Headspace volatiles detected are shown in GC-MS chromatograms for each *Ipomoea* sp. All experimentally tested compounds were detected in the headspace odor.

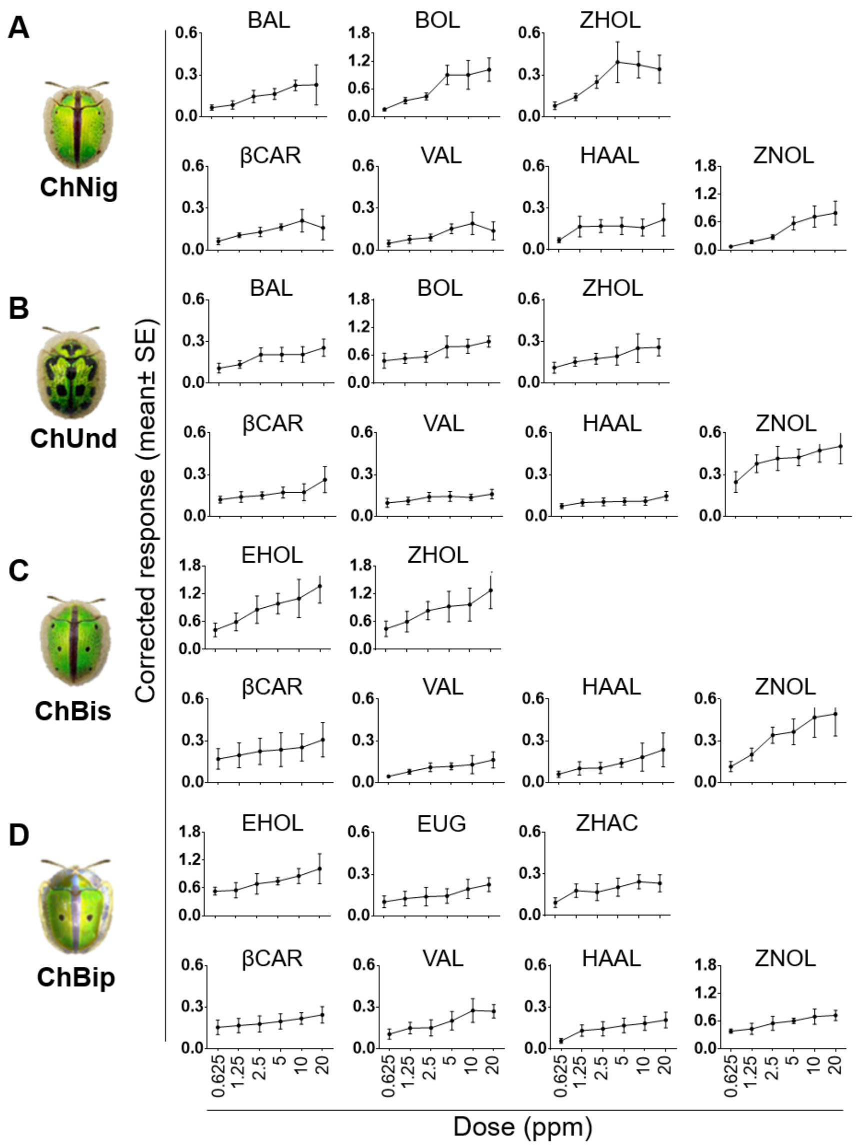


**Figure S6: Experimentally tested attractants, repellents, and neutrals are EAG-active.** Electrophysiological responses were observed in antennae of (**A**) *C. nigropunctata*, (**B**) *C. undecimnotata*, (**C**) *C. bistrimaculata*, and (**D**) *C. bipunctata* towards all the experimental candidate compounds, suggesting that they have olfactory receptors for all the compounds. Increasing EAG response was observed with increasing doses of compounds. The data shown in the figure is the blank subtracted response. Blank in all cases was DCM.

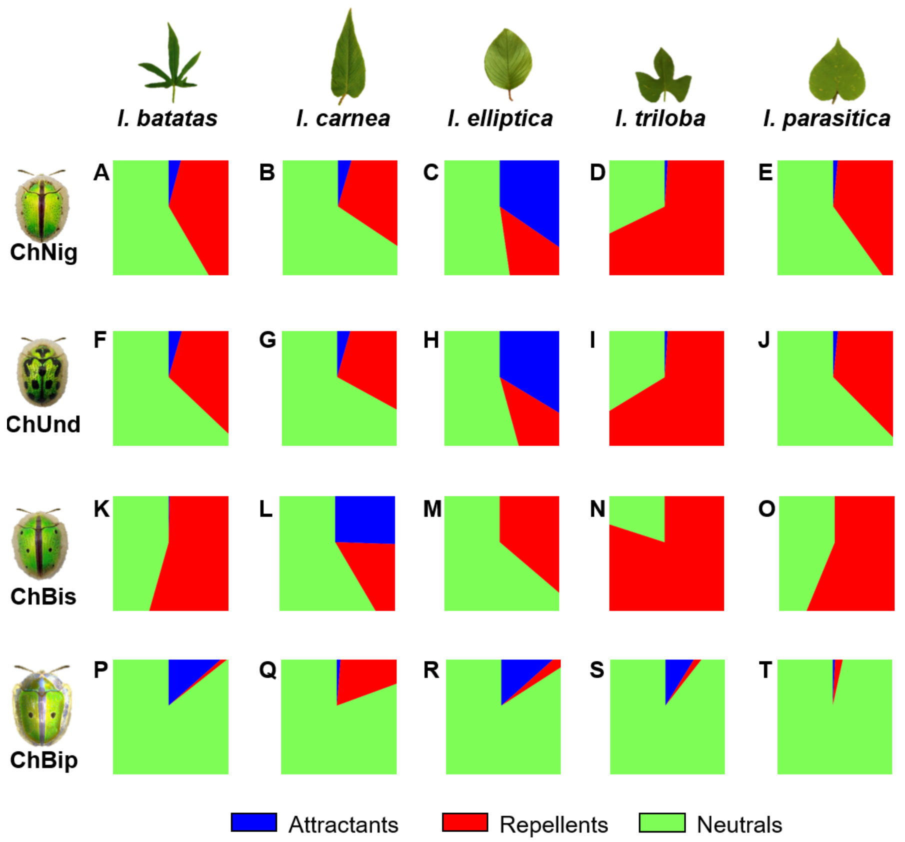


**Figure S7: Each *Ipomoea* sp.'s odor blend has a signature proportion of attractants, repellents, and neutrals, which differs for every *Chiridopsis* sp.** For every *Chiridopsis* sp., the concentration of each attractant (blue), repellent (red), and neutral compound (green) (Fig 4F) was multiplied by the beetle's standardized regression coefficient for that compound. Resulting values were plotted as a pie diagram for (**A-E**) *C. nigropunctata*, (**F-J**) *C. undecimnotata*, (**K-O**) *C. bistrimaculata,* and (**P-T**) *C. bipunctata*, resulting in a different pie diagram for each beetle-plant pair. Pie diagrams were plotted as squares instead of circles for ease of using them to pixelate leaf shapes (Fig 6). Each pie diagram represents a plant's odor as perceived by a particular beetle.

**Table S1: Volatile compounds detected in *Ipomoea* spp.** Concentration of 29 volatile compounds identified from *I. batatas, I. carnea, I. elliptica, I. triloba* and *I. parasitica* (nmol/ g leaf tissue, mean± SE, n= 6). Reported concentrations are estimated by normalizing relative to nonyl acetate internal standard. nd= Not detected in analysis.

| Compound | | | Kovat's retention index | | Concentration in *Ipomoea* spp. (nmol/ g, mean± SE) | | | | |
|---|---|---|---|---|---|---|---|---|---|
| **Class** | **Name** | **Abbreviation** | **Experimentally determined** | **Reported on NIST-MS library** | ***I. batatas*** | ***I. carnea*** | ***I. elliptica*** | ***I. triloba*** | ***I. parasitica*** |
| Alcohols | Benzyl alcohol | BOL | 1043 | 1036 | 4.51 ± 0.42 | 2.87 ± 0.48 | nd | nd | 2.51 ± 0.26 |
| | Phenylethyl Alcohol | PEA | 1119 | 1116 | 44.81 ± 5.76 | 3.96 ± 0.36 | nd | nd | nd |
| | β-Phenoxyethyl alcohol | βPEA | 1228 | 1225 | 2.6 ± 0.19 | nd | nd | nd | nd |
| | (E)-Hex-3-en-1-ol | EHOL | 866 | 852 | 8.29 ± 0.3 | nd | nd | nd | nd |
| | (Z)-Hex-3-en-1-ol | ZHOL | 861 | 861/857 | 64.36 ± 5.33 | 35.91 ± 7.65 | 30.63 ± 4.33 | 70.06 ± 9.93 | 111.25 ± 15.57 |
| Aldehydes | Benzeneacetaldehyde | BAL | 1050 | 1045 | 8.29 ± 0.93 | 11.97 ± 2.59 | 19.26 ± 2.24 | 8.52 ± 1.04 | 9.02 ± 1.04 |
| | 3-Hexenal | HAL | 803 | 810 | 15.61 ± 0.35 | nd | 25.66 ± 3.02 | nd | nd |
| Oxygenated terpenes | Camphol | CAM | 1172 | 1167 | 0.33 ± 0.04 | 0.43 ± 0.05 | 0.36 ± 0.03 | 0.39 ± 0.02 | 0.28 ± 0.02 |
| | Germacrene D-4-ol | GOL | 1494 | 1574 | 3.86 ± 0.42 | 4.13 ± 0.83 | 11.78 ± 3.25 | nd | 11.42 ± 1.37 |
| Sesquiterpenes | γ-Elemene | γELE | 1434 | 1434 | 136.25 ± 19.8 | nd | 15.34 ± 3.56 | 258.81 ± 22.92 | 260.19 ± 25.89 |
| | δ-Elemene | δELE | 1344 | 1338 | nd | 83.11 ± 14.47 | nd | nd | nd |
| | α-Copaene | αCOP | 1382 | 1376 | 23.29 ± 2.49 | 2.08 ± 0.39 | 45.89 ± 12.39 | 42.87 ± 3.8 | 11.17 ± 1.42 |
| | β-Cubebene | βCUB | 1396 | 1389 | 9.66 ± 1.56 | 1.5 ± 0.1 | 1.65 ± 0.38 | 4.65 ± 0.31 | 7.07 ± 1.32 |
| | β-Elemene | βELE | 1397 | 1391 | 115.24 ± 12.84 | 16.85 ± 1.54 | 19.91 ± 4.95 | 27.28 ± 2.07 | 126.02 ± 14.72 |
| | β-Caryophyllene | βCAR | 1416 | 1419 | 542.51 ± 57.21 | 206.11 ± 24.91 | 123.54 ± 22.02 | 194.67 ± 20.45 | 912.03 ± 112.38 |
| | β-Copaene | βCOP | 1451 | 1432 | 29.47 ± 3.03 | 24.97 ± 5 | 21.3 ± 8.13 | 20.15 ± 2.24 | 77.92 ± 8.07 |
| | α-Bergamotene | αBER | 1423 | 1415 | 68.48 ± 5.29 | 17.94 ± 3.95 | 36.01 ± 13.06 | 2.69 ± 0.38 | 8.11 ± 0.84 |
| | Isogermacrene D | IGD | 1427 | 1448 | 5.7 ± 0.4 | 12.61 ± 2.45 | 12.23 ± 4.4 | 8.94 ± 0.91 | 32.54 ± 3.3 |
| | Humulene | HUM | 1434 | 1454 | 80.5 ± 7.95 | 38.03 ± 4.4 | 129.67 ± 22.63 | 37.83 ± 3.92 | 182.4 ± 21.14 |
| | epi-β-Caryophyllene | eβC | 1439 | 1440 | nd | nd | 134.71 ± 55.44 | nd | nd |
| | Germacrene D | GD | 1451 | 1481 | 543.49 ± 52.23 | 463.2 ± 93.22 | 388.06 ± 148.05 | 367.28 ± 42.24 | 1433.55 ± 145.45 |
| | Bicyclogermacrene | BCG | 1457 | 1495 | 175.6 ± 24.68 | 21.29 ± 4.17 | 80.66 ± 29.71 | 409.49 ± 35.11 | 439.63 ± 48.17 |
| | α-Farnesene | αFAR | 1460 | 1508 | 14.24 ± 0.78 | 32.44 ± 7.42 | nd | 27.14 ± 2.66 | 11.68 ± 3.26 |
| | β-Cadinene | βCAD | 1469 | 1518 | nd | nd | 15.68 ± 5.88 | nd | nd |
| | δ-Cadinene | δCAD | 1469 | 1524 | 13.69 ± 0.94 | 2.46 ± 0.37 | 49.37 ± 29.69 | 18.25 ± 1.27 | 1.96 ± 0.36 |
| | Germacrene B | GB | 1486 | 1557 | 3.48 ± 0.28 | 3.67 ± 0.45 | nd | nd | nd |
| Other | 1,3-Di-tertiary-butylbenzene | DTB | 1259 | 1281 | 13.27 ± 2.34 | 13.29 ± 1.1 | 19.46 ± 0.77 | 21.6 ± 1.46 | 11.78 ± 1.01 |
| | (Z)-3-Hexenyl Acetate | ZHAC | 1012 | 1005 | 1.44 ± 0.26 | 5.55 ± 1.28 | 2.35 ± 0.44 | nd | 4.61 ± 0.66 |
| | Eugenol | EUG | 1365 | 1357 | 4.81 ± 0.52 | 1.14 ± 0.1 | nd | nd | nd |
| Total concentration | | | | | 1933.8 ± 85.71 | 1005.51 ± 98.65 | 1183.51 ± 168.17 | 1520.62 ± 64.12 | 3655.13 ± 194.37 |

**Table S2: Factor loading, percentage contribution, and squared cosines of variables (volatile compounds) on the first two principal components of PCA.** For each variable, values in bold correspond to the factor for which the squared cosine is the largest among all the extracted principal components.

| Volatile compound | Factor loadings | | Contribution (%) | | Squared cosines | |
|---|---|---|---|---|---|---|
| | PC1 | PC2 | PC1 | PC2 | PC1 | PC2 |
| HAL | -0.253 | 0.218 | 0.799 | 0.698 | 0.064 | 0.047 |
| EHOL | 0.395 | 0.838 | 1.940 | 10.352 | 0.156 | **0.702** |
| ZHOL | 0.677 | -0.287 | 5.701 | 1.215 | **0.458** | 0.082 |
| ZHAC | 0.166 | -0.184 | 0.343 | 0.500 | 0.028 | 0.034 |
| BOL | 0.580 | 0.639 | 4.190 | 6.020 | 0.337 | **0.408** |
| BAL | -0.407 | -0.260 | 2.061 | 0.995 | 0.166 | 0.067 |
| PEA | 0.389 | 0.844 | 1.880 | 10.517 | 0.151 | **0.713** |
| CAM | -0.539 | 0.150 | 3.616 | 0.331 | **0.291** | 0.022 |
| βPEA | 0.366 | 0.822 | 1.671 | 9.972 | 0.134 | **0.676** |
| DTB | -0.558 | -0.228 | 3.868 | 0.767 | **0.311** | 0.052 |
| γELE | 0.641 | -0.329 | 5.109 | 1.601 | **0.411** | 0.109 |
| δELE | -0.269 | 0.184 | 0.901 | 0.502 | 0.072 | 0.034 |
| EUG | 0.359 | 0.893 | 1.601 | 11.768 | 0.129 | **0.798** |
| αCOP | -0.325 | -0.160 | 1.316 | 0.380 | 0.106 | 0.026 |
| βCUB | 0.798 | 0.304 | 7.919 | 1.366 | **0.637** | 0.093 |
| βELE | 0.965 | 0.096 | 11.581 | 0.137 | **0.931** | 0.009 |
| βCAR | 0.962 | -0.149 | 11.524 | 0.327 | **0.926** | 0.022 |
| βCOP | 0.813 | -0.480 | 8.212 | 3.400 | **0.660** | 0.231 |
| αBER | 0.164 | 0.631 | 0.335 | 5.872 | 0.027 | 0.398 |
| IGD | 0.577 | -0.688 | 4.139 | 6.982 | 0.333 | **0.473** |
| HUM | 0.518 | -0.461 | 3.333 | 3.136 | 0.268 | 0.213 |
| eβC | -0.306 | -0.295 | 1.162 | 1.285 | 0.093 | 0.087 |
| GD | 0.813 | -0.478 | 8.217 | 3.366 | **0.661** | 0.228 |
| BCG | 0.603 | -0.492 | 4.527 | 3.564 | **0.364** | 0.242 |
| αFAR | -0.108 | 0.207 | 0.145 | 0.634 | 0.012 | 0.043 |
| βCAD | -0.348 | -0.251 | 1.506 | 0.926 | 0.121 | 0.063 |
| δCAD | -0.349 | -0.025 | 1.515 | 0.009 | 0.122 | 0.001 |
| GB | 0.079 | 0.807 | 0.079 | 9.614 | 0.006 | **0.652** |
| GOL | 0.255 | -0.505 | 0.807 | 3.765 | 0.065 | 0.255 |

**Table S3:** Correlation coefficients and *p* values for volatile compounds (independent variables) and *Chiridopsis* feeding preference (dependent variables) on the first two principal components. *p* values in bold are significant after sequential Bonferroni correction.

| Variable | Correlation | | *p* value | |
|---|---|---|---|---|
| | **Component 1** | **Component 2** | **Component 1** | **Component 2** |
| Volatile compound | | | | |
| HAL | 0.730 | 0.459 | **<0.0001** | 0.0107 |
| EHOL | -0.291 | 0.694 | 0.1187 | **<0.0001** |
| ZHOL | -0.585 | 0.314 | **0.0007** | 0.0911 |
| ZHAC | -0.214 | -0.536 | 0.2562 | **0.0023** |
| BOL | -0.645 | 0.180 | **0.0001** | 0.3412 |
| BAL | 0.710 | -0.030 | **<0.0001** | 0.875 |
| PEA | -0.320 | 0.646 | 0.0847 | **0.0001** |
| CAM | 0.135 | -0.460 | 0.4769 | 0.0105 |
| βPEA | -0.276 | 0.658 | 0.1399 | **0.0001** |
| DTB | 0.488 | -0.018 | 0.0072 | 0.9262 |
| γELE | -0.535 | 0.472 | **0.0028** | 0.0097 |
| δELE | -0.224 | -0.774 | 0.2428 | **<0.0001** |
| EUG | -0.363 | 0.542 | 0.0529 | **0.0024** |
| αCOP | 0.592 | 0.424 | **0.0007** | 0.0219 |
| βCUB | -0.517 | 0.779 | 0.0041 | **<0.0001** |
| βELE | -0.563 | 0.743 | **0.0015** | **<0.0001** |
| βCAR | -0.675 | 0.494 | **<0.0001** | 0.0065 |
| βCOP | -0.439 | 0.242 | 0.0172 | 0.2059 |
| αBER | 0.178 | 0.607 | 0.3556 | **0.0005** |
| IGD | -0.183 | -0.044 | 0.3420 | 0.8207 |
| HUM | 0.153 | 0.458 | 0.4281 | 0.0125 |
| eβC | 0.711 | 0.089 | **<0.0001** | 0.6462 |
| GD | -0.443 | 0.235 | 0.0161 | 0.2198 |
| BCG | -0.418 | 0.415 | 0.0240 | 0.0252 |
| αFAR | -0.590 | -0.532 | **0.0008** | **0.0030** |
| βCAD | 0.760 | 0.129 | **<0.0001** | 0.5048 |
| δCAD | 0.496 | 0.195 | 0.0062 | 0.3107 |
| GB | -0.428 | -0.174 | 0.0205 | 0.3667 |
| GOL | 0.420 | 0.168 | 0.0233 | 0.3837 |
| Feeding preference | | | | |
| ChNig | 0.935 | 0.092 | **<0.0001** | 0.6350 |
| ChBis | 0.249 | -0.652 | 0.1927 | **0.0001** |
| ChBip | 0.323 | 0.613 | 0.0874 | **0.0004** |
| ChUnd | 0.910 | 0.232 | **<0.0001** | 0.2259 |

**Table S4**: Intercept ($\beta_0$) and coefficients ($\beta_i$) of multiple regression ($y = \beta_0 + \sum \beta_i x_i$) between feeding preference ($y$) and volatile metabolites ($x_i$). The goodness of fit ($R^2$) values are provided at the bottom.

| Variable | ChNig | ChBis | ChBip | ChUnd |
|---|---|---|---|---|
| Intercept | 22.6856 | 28.9020 | 19.3022 | 20.7306 |
| HAL | 0.5772 | 0.0323 | 0.2064 | 0.5252 |
| EHOL | -0.2771 | -0.2919 | 0.6116 | 0.0661 |
| ZHOL | -0.0724 | -0.0535 | -0.0192 | -0.0626 |
| ZHAC | -0.0634 | 1.1222 | -0.7937 | -0.2168 |
| BOL | -1.6462 | 0.6528 | -0.2087 | -1.1151 |
| BAL | 0.8351 | 0.2169 | 0.1559 | 0.6979 |
| PEA | -0.0521 | -0.0307 | 0.0978 | 0.0080 |
| CAM | -10.1705 | 13.3538 | -7.9009 | -11.3822 |
| βPEA | -0.9370 | -0.9071 | 1.7738 | 0.0807 |
| DTB | 0.2743 | -0.1925 | 0.1528 | 0.2056 |
| γELE | -0.0326 | -0.0294 | 0.0003 | -0.0265 |
| δELE | -0.0541 | 0.0817 | -0.0473 | -0.0529 |
| EUG | -0.6912 | -0.0242 | 0.8659 | -0.0697 |
| αCOP | 0.1473 | -0.0666 | 0.0837 | 0.1333 |
| βCUB | -0.5257 | -0.5211 | 0.3236 | -0.2458 |
| βELE | -0.0301 | -0.0333 | 0.0113 | -0.0148 |
| βCAR | -0.0070 | -0.0044 | -0.0011 | -0.0050 |
| βCOP | -0.0544 | -0.0245 | -0.0550 | -0.0549 |
| αBER | 0.0767 | -0.0021 | 0.0613 | 0.0867 |
| IGD | -0.0123 | 0.0108 | -0.1886 | -0.0720 |
| HUM | 0.0631 | -0.0121 | 0.0039 | 0.0513 |
| eβC | 0.0591 | 0.0084 | 0.0108 | 0.0488 |
| GD | -0.0031 | -0.0013 | -0.0031 | -0.0031 |
| BCG | -0.0175 | -0.0189 | -0.0015 | -0.0153 |
| αFAR | -0.3527 | 0.0426 | -0.1091 | -0.3087 |
| βCAD | 0.5660 | 0.0374 | 0.0414 | 0.4371 |
| δCAD | 0.0842 | -0.0129 | 0.0250 | 0.0700 |
| GB | -1.3983 | 1.1767 | -0.0593 | -0.9007 |
| GOL | 0.7593 | 0.1373 | 0.0011 | 0.6005 |
| $R^2$ | 0.9186 | 0.6900 | 0.6399 | 0.9093 |

**Table S5**: Standardized coefficients and *p* values for the null hypothesis that coefficients are not significantly different from zero. Numbers in brackets denote standard error. Significant *p* values (≤ 0.005) after sequential Bonferroni correction are shown in bold.

| Variable | ChNig | | ChBis | | ChBip | | ChUnd | |
|---|---|---|---|---|---|---|---|---|
| | Coefficient | *p* | Coefficient | *p* | Coefficient | *p* | Coefficient | *p* |
| HAL | 0.1552 (0.0253) | **<0.0001** | 0.0164 (0.0356) | 0.6581 | 0.1184 (0.0343) | **0.0006** | 0.1739 (0.0276) | **<0.0001** |
| EHOL | -0.0220 (0.0083) | 0.0084 | -0.0437 (0.0130) | **0.0008** | 0.1034 (0.0162) | **<0.0001** | 0.0065 (0.0083) | 0.4428 |
| ZHOL | -0.0633 (0.0169) | **0.0002** | -0.0882 (0.0303) | **0.0036** | -0.0357 (0.0209) | 0.0866 | -0.0674 (0.0197) | **0.0007** |
| ZHAC | -0.0038 (0.0214) | 0.8694 | 0.1266 (0.0476) | 0.0078 | -0.1012 (0.0319) | **0.0016** | -0.0160 (0.0231) | 0.4990 |
| BOL | -0.0696 (0.0145) | **<0.0001** | 0.0521 (0.0274) | 0.0574 | -0.0188 (0.0379) | 0.6325 | -0.0581 (0.0168) | **0.0006** |
| BAL | 0.1138 (0.0288) | **0.0001** | 0.0558 (0.0663) | 0.4073 | 0.0453 (0.0401) | 0.2618 | 0.1172 (0.0296) | **0.0001** |
| PEA | -0.0236 (0.0068) | **0.0006** | -0.0263 (0.0112) | 0.0186 | 0.0945 (0.0197) | **<0.0001** | 0.0044 (0.0082) | 0.5984 |
| CAM | -0.0232 (0.0262) | 0.3817 | 0.0575 (0.0511) | 0.2636 | -0.0385 (0.0360) | 0.2893 | -0.0320 (0.0300) | 0.2897 |
| βPEA | -0.0238 (0.0097) | 0.0145 | -0.0435 (0.0140) | **0.0019** | 0.0960 (0.0188) | **<0.0001** | 0.0025 (0.0101) | 0.8141 |
| DTB | 0.0335 (0.0219) | 0.1259 | -0.0443 (0.0202) | 0.0279 | 0.0398 (0.0328) | 0.2271 | 0.0309 (0.0234) | 0.1873 |
| γELE | -0.0874 (0.0144) | **<0.0001** | -0.1488 (0.0163) | **<0.0001** | 0.0016 (0.0353) | 0.9663 | -0.0877 (0.0156) | **<0.0001** |
| δELE | -0.0492 (0.0150) | **0.0011** | 0.1404 (0.0472) | **0.0030** | -0.0919 (0.0244) | **0.0002** | -0.0593 (0.0164) | **0.0003** |
| EUG | -0.0324 (0.0070) | 0.0000 | -0.0021 (0.0151) | 0.8957 | 0.0865 (0.0223) | **0.0001** | -0.0040 (0.0090) | 0.6665 |
| αCOP | 0.0778 (0.0305) | 0.0108 | -0.0664 (0.0253) | 0.0088 | 0.0944 (0.0340) | **0.0055** | 0.0868 (0.0358) | 0.0152 |
| βCUB | -0.0462 (0.0141) | **0.0011** | -0.0864 (0.0174) | **<0.0001** | 0.0606 (0.0335) | 0.0702 | -0.0266 (0.0176) | 0.1310 |
| βELE | -0.0350 (0.0072) | **<0.0001** | -0.0733 (0.0121) | **<0.0001** | 0.0280 (0.0233) | 0.2327 | -0.0213 (0.0088) | 0.0150 |
| βCAR | -0.0472 (0.0174) | 0.0066 | -0.0555 (0.0225) | 0.0138 | -0.0164 (0.0211) | 0.4433 | -0.0417 (0.0153) | 0.0064 |
| βCOP | -0.0277 (0.0201) | 0.1694 | -0.0235 (0.0231) | 0.3143 | -0.0597 (0.0232) | 0.0100 | -0.0344 (0.0211) | 0.1030 |
| αBER | 0.0509 (0.0361) | 0.1590 | -0.0027 (0.0361) | 0.9458 | 0.0868 (0.0321) | 0.0069 | 0.0709 (0.0358) | 0.0475 |
| IGD | -0.0028 (0.0265) | 0.9222 | 0.0047 (0.0280) | 0.8766 | -0.0922 (0.0263) | **0.0005** | -0.0203 (0.0273) | 0.4653 |
| HUM | 0.0873 (0.0365) | 0.0168 | -0.0315 (0.0229) | 0.1706 | 0.0116 (0.0405) | 0.7865 | 0.0875 (0.0348) | 0.0120 |
| eβC | 0.1157 (0.0302) | **0.0001** | 0.0310 (0.0577) | 0.6035 | 0.0452 (0.0306) | 0.1403 | 0.1177 (0.0307) | **0.0001** |
| GD | -0.0287 (0.0199) | 0.1500 | -0.0233 (0.0230) | 0.3170 | -0.0611 (0.0234) | 0.0091 | -0.0357 (0.0209) | 0.0881 |
| BCG | -0.0703 (0.0195) | **0.0003** | -0.1439 (0.0168) | **<0.0001** | -0.0132 (0.0348) | 0.7179 | -0.0758 (0.0196) | **0.0001** |
| αFAR | -0.1250 (0.0244) | **<0.0001** | 0.0285 (0.0701) | 0.6973 | -0.0825 (0.0328) | 0.0119 | -0.1348 (0.0255) | **<0.0001** |
| βCAD | 0.1231 (0.0272) | **<0.0001** | 0.0153 (0.0675) | 0.8314 | 0.0192 (0.0309) | 0.5460 | 0.1171 (0.0310) | **0.0002** |
| δCAD | 0.0738 (0.0344) | 0.0315 | -0.0213 (0.0233) | 0.3658 | 0.0467 (0.0521) | 0.3769 | 0.0756 (0.0390) | 0.0519 |
| GB | -0.0609 (0.0126) | **<0.0001** | 0.0968 (0.0268) | **0.0003** | -0.0055 (0.0339) | 0.8804 | -0.0484 (0.0149) | **0.0012** |
| GOL | 0.1044 (0.0231) | **<0.0001** | 0.0356 (0.0382) | 0.3571 | 0.0003 (0.0400) | 0.9941 | 0.1017 (0.0262) | **0.0001** |
